# CRISPR screens in 3D assembloids reveal disease genes associated with human interneuron development

**DOI:** 10.1101/2022.09.06.506845

**Authors:** Xiangling Meng, David Yao, Xiaoyu Chen, Kevin W. Kelley, Noah Reis, Mayuri Vijay Thete, Shravanti Kulkarni, Michael C. Bassik, Sergiu P. Pașca

## Abstract

The assembly of cortical circuits involves the generation and migration of cortical interneurons from the ventral to the dorsal forebrain, which has been challenging to study in humans as these processes take place at inaccessible stages of late gestation and early postnatal development. Autism spectrum disorder (ASD) and other neurodevelopmental disorders (NDDs) have been associated with abnormal cortical interneuron development, but which of the hundreds of NDD genes impact interneuron generation and migration into circuits and how they mediate these effects remain unknown. We previously developed a stem cell-based platform to study human cortical interneurons in self-organizing organoids resembling the ventral forebrain and their migration using forebrain assembloids. Here, we integrate assembloid technology with CRISPR screening to systematically investigate the involvement of 425 NDD genes in human interneuron development. The first screen aimed at interneuron generation revealed 13 candidate genes, including the RNA-binding protein *CSDE1* and the canonical TGFβ signaling activator *SMAD4*. Then, we ran an interneuron migration screen in ∼1,000 forebrain assembloids that identified 33 candidate genes, including cytoskeleton-related genes and, notably, the endoplasmic reticulum (ER)-related gene *LNPK*. Interestingly, we discovered that, during interneuron migration, the ER is displaced along the leading neuronal branch prior to nuclear translocation. Deletion of *LNPK* interfered with this ER displacement and resulted in reduced interneuron saltation length, indicating a critical role for the ER in this migratory process. Taken together, these results highlight how this versatile CRISPR-assembloid platform can be used to systematically map disease genes onto early stages of human neural development and to reveal novel mechanisms regulating interneuron development.

## Introduction

Defects in the development and function of cortical GABAergic interneurons have been implicated in ASD and other NDDs^1,2^. This is primarily based on the high prevalence of seizures and epilepsy in patients^2-5^ and neuroimaging studies that revealed changes in interneuron density and morphology in postmortem ASD tissue^6,7^. Concurrently, whole-exome sequencing and genome-wide association studies over the past decade have uncovered hundreds of disease-related genes for ASD and other NDDs^8-11^, including several GABA receptor subunit genes related to inhibitory neural transmission^12,13^. It has been, however, challenging to map the role of these disease genes onto specific stages of the protracted human interneuron development.

Cortical GABAergic interneurons are generated in the ventral forebrain and they subsequently migrate long-distances towards the dorsal forebrain to integrate into circuits^14-18^. In humans this process takes place in later stages of gestation and, unlike in rodents, continues into the postnatal period^19,20^. This protracted interneuron development is thought to contribute to increased complexity and cognitive abilities in the gyrencephalic brain^20^. To model stages of human interneuron development, we and others have developed self-organizing organoids resembling the ventral forebrain – human subpallial organoids (hSO), from human induced pluripotent stem (hiPS) cells^21-23^. These organoids can be integrated with human cortical organoids (hCO) to form human forebrain assembloids (hFA) and model interneuron migration and circuit assembly with glutamatergic neurons^21-23^. These stem cell-based 3D cellular models of development and disease have either focused on studying the role of single genes in interneuron generation or migration^21,24-26^ or have leveraged screens to study the role of groups of genes in early stages of neural proliferation^27^. Mapping in parallel the impact of hundreds of disease genes onto multiple stages of interneuron development, including their migration into cortical circuits at later stages of gestation, has not been achieved.

Here, we coupled human organoid and assembloid technologies with CRISPR screening to map the roles of a group of subpallium-expressed NDD genes onto interneuron generation and migration into cortical circuits. We discovered that 46 out the of 425 disease genes (∼11%) interfere with interneuron development. Notably, we discovered that loss of the ER-shaping protein LNPK disrupts interneuron migration, highlighting a previously unappreciated role of ER dynamics in interneuron migration towards the cerebral cortex..

## Results

### Pooled loss-of-function CRISPR screens reveal NDD genes related to interneuron generation and migration

To determine which disease genes interfere with cortical interneuron development, we first generated a knock-in hiPS cell line that stably expresses Cas9 and that can be turned off with doxycycline (Tet-Off). To achieve this, we inserted CAG::Cas9 into the *AAVS1* locus (**Supplementary Fig. 1a**), generated a clonal CAG::Cas9 hiPS cell line (**Supplementary Fig. 1b–d**), and confirmed the expression of Cas9 (**Supplementary Fig. 1e**) as well as the editing efficiency by infecting the cells with a lentivirus harboring GFP and a sgRNA against GFP. Using this strategy, we observed an efficient reduction in the proportion of GFP^+^ cells within 9 days of Cas9 expression (**Supplementary Fig. 1f**). To facilitate isolation of ventral forebrain interneuron lineages, we further modified the CAG::Cas9 line to stably express GFP under the control of a previously validated enhancer Dlxi1/2b that labels medial ganglionic eminence (MGE)-derived cells^21,28^. When we differentiated this CAG::Cas9;Dlxi1/2b::eGFP hiPS cell line into hSO, we detected extensive expression of GFP after 30 days of differentiation. When we assembled hSO with hCO derived from an unlabeled hiPS cell line for 30 days, we observed a notable fraction of Dlxi1/2b::eGFP^+^ cells that migrated into the cortical side of the hFA (**Supplementary Fig. 1g**).

To screen for genes associated with ASD and other NDDs, we compiled a list of 611 genes (**Supplementary Table 1**) based on genetic studies of neurodevelopmental disorder patients^8-11^ and the Simons Foundation Autism Research initiative (SFARI) database (score 1, 2, and syndromic) (**Supplementary Fig. 2a**). After excluding genes that are not expressed in hSO Dlxi1/2::eGFP^+^ cells or human embryonic ganglionic eminences (GE), we ended up with 425 NDD genes (**Fig. 1a**, 354 ASD genes and 71 genes associated with other NDDs. **Supplementary Fig.2b** and **Supplementary Table 2**). These genes showed higher expression during prenatal than postnatal stages (**Supplementary Fig. 2c**), consistent with their roles in neural development. We then synthesized a sgRNA library containing 5 sgRNAs per candidate gene together with 13 genes with known function in interneurons and ‘safe’ harbor-targeting negative controls (218 sgRNAs), and transduced it into CAG::Cas9;Dlxi1/2b::eGFP hiPS cells (1,000X coverage per sgRNA). The lentivirus encoding the sgRNA library also express mCherry, and we used fluorescence-activated cell sorting (FACS) to select transduced mCherry^+^ cells for differentiation into hSO 6 days later (**Fig. 1b**). To verify that our platform can detect genes that perturb early stages of neurodevelopment, such as proliferation, we performed the experiment without doxycycline and looked at proliferation related NDD genes. Indeed, sgRNAs targeting *TSC2, TSC1*, and *PTEN*, which are known to promote neural proliferation when lost^29-31^, were enriched in hSO at 19 days of differentiation compared with hiPS cells (**Supplementary Fig. 3a**).

**Figure 1:**
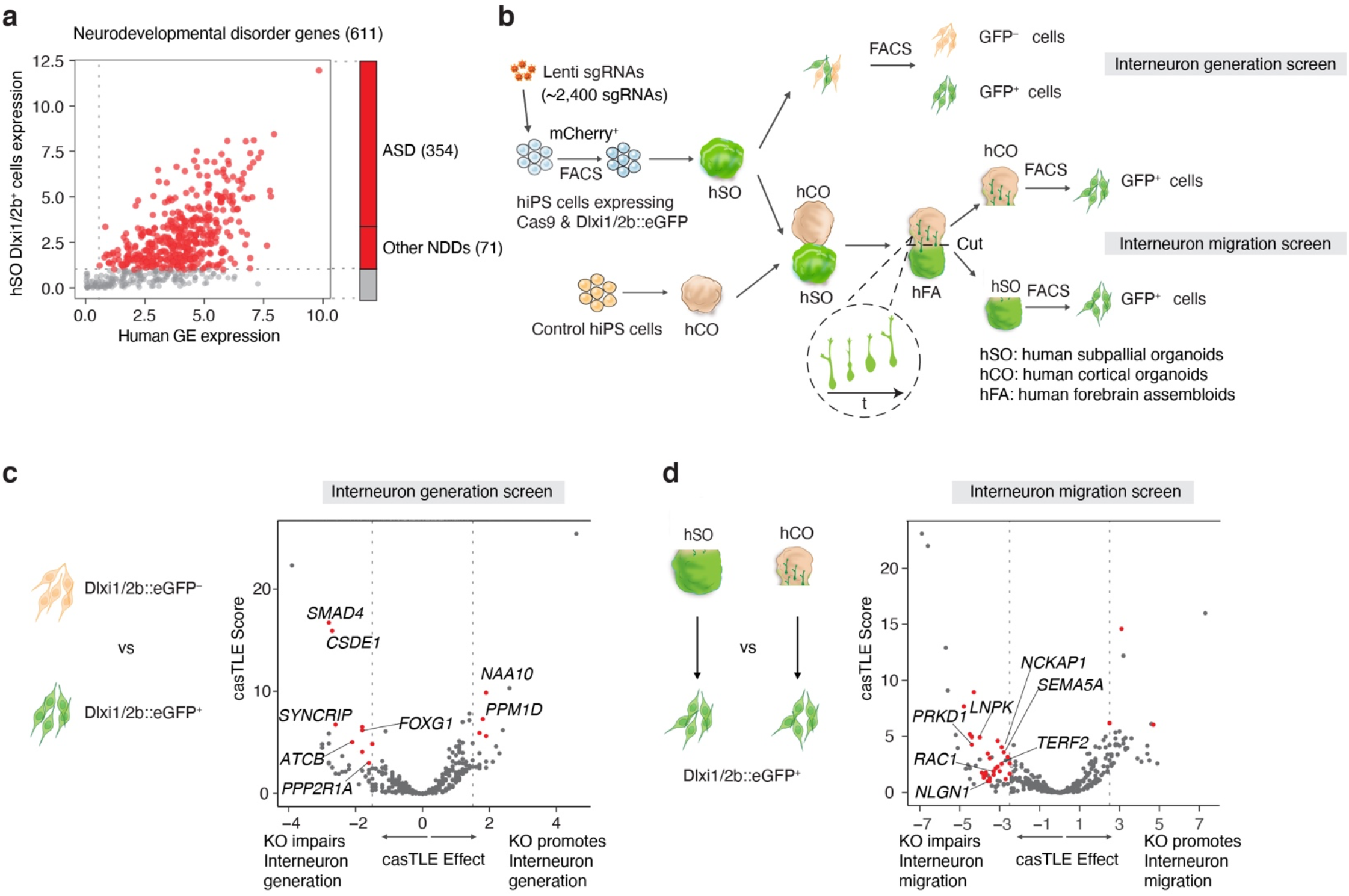
CRISPR screens of NDD genes reveal regulators of human interneuron generation and migration. **a**, A total of 425 genes (labeled in red) out of 611 NDD genes showing expression in hSO Dlxi1/2b::eGFP^+^ cells and human PCW 8-9 ganglionic eminences were included in the CRISPR screens. Among these, 354 genes are associated with ASD and 71 genes are associated with other NDDs. **b**, Schematic describing the interneuron generation and migration CRISPR screens. Inset highlights how hSO-derived interneurons migrate into the hCO side of the hFA. hSO: human subpallial organoids; hCO: human cortical organoids; hFA: human forebrain assembloids. **c**, Volcano plot of the CasTLE-estimated maximum gene perturbation effect size and associated CasTLE score for the interneuron generation screen. Genes with at least 2 sgRNAs having effects ≤ –1.5 or ≥ +1.5, approximately 2x the standard deviation (SD) of negative controls were selected as candidates (red), with labels for cell cycle genes and genes that were subsequently validated. **d**, Volcano plot of the CasTLE-estimated maximum gene perturbation effect size and associated CasTLE score for the interneuron migration screen. Genes with at least 2 sgRNAs having effects ≤ –2.5 or ≥ +2.5, approximately 2x the SD of negative controls were selected as candidates (red), with labels for cytoskeleton and cell migration genes and genes that were subsequently validated

To identify genes that affect interneuron generation, we collected hSO at 44 days of differentiation and used FACS to separate the Dlxi1/2b::eGFP^+^ cells (**Fig. 1b**). We evaluated the relative enrichment of sgRNAs for each targeted gene when comparing the eGFP^+^ and eGFP^−^ populations, and we identified 13 genes whose sgRNAs have concordant effects that significantly deviate from the negative control sgRNA distribution (**Fig. 1c** and **Supplementary Table 3**). This list included 5 cell cycle-related genes, including *PPP2R1A, PPM1D*, as well as *FOXG1*, which is critical for forebrain interneuron development^32,33^ and the loss of which results in NDDs^34,35^.

To prioritize ASD and NDD genes that regulate interneuron migration, we next created hFA by integrating unperturbed hiPS cell-derived hCO at day 60 and CRISPR-perturbed hiPS cell-derived hSO at 45 days of differentiation (**Fig.1b**). We generated ∼1,000 hFA to achieve an estimated coverage of 500 migrated cells per sgRNA. We allowed interneurons from hSO to migrate into hCO for 30 days; we then separated the two sides of the hFA and used FACS to isolate Dlxi1/2b::eGFP^+^ cells from the hSO part and migrated Dlxi1/2b::eGFP^+^ cells from the hCO part (**Fig. 1b**). We evaluated the relative enrichment of sgRNAs for each targeted gene in migrated and non-migrated Dlxi1/2b::eGFP^+^ cells, and we identified 33 candidate genes affecting interneuron migration (**Fig. 1d** and **Supplementary Table 4**). Five of these genes were related to cytoskeleton organization and cell migration, including *NCKAP1, SEMA5A, PRKD1*, as well as *RAC1*– a major regulator of cytoskeleton dynamics^36^. This is consistent with previous studies suggesting that functional repression of Rac1 in mice impairs neuronal migration by affecting the leading process of migrating interneurons^37-39^.

### Loss of *SMAD4* and *CSDE1* impairs subpallium differentiation

To confirm the role of the candidate genes in subpallium development, we decided to validate the top hits from the interneuron generation screen: *SMAD4, CSDE1*, and *SYNCRIP* (**Fig. 1c** and **Supplementary Fig. 4a**). We engineered the CAG::Cas9;Dlxi1/2b::eGFP hiPS cell line by nucleofecting three sgRNAs per gene targeting each of these gene hits and generated KO hiPS cell pools for each candidate (**Fig. 2a** and **Supplementary Fig. 4b**). Both hiPS cells and hSO at 40 days of differentiation showed efficient deletion of the corresponding genes (**Supplementary Fig. 4c, d**). We found that KO cell pool-derived hSO at 40 days of differentiation contained more than 96% of sequenced alleles with partial deletion (**Supplementary Fig. 4e–g**). At 45 days of differentiation, we examined hSO by flow cytometry and found that *SMAD4* and *CSDE1* KO hSO (****P < 0.0001), but not *SYNCRIP* KO hSO (P = 0.87), generated reduced proportions of

Dlxi1/2b::eGFP^+^ cells compared with the Cas9-control (Cas9-CTL, **Fig 2b, c**). Furthermore, we found that *SMAD4* and *CSDE1* KO hSO were smaller in size than Cas9-CTL at multiple *in vitro* differentiation stages (**Fig. 2d, e**, P < 0.01). Several studies have suggested that both *SMAD4* and *CSDE1* are involved in neural differentiation^40-42^, so we next examined the expression of transcription factors that regulate subpallium development. Consistent with the reduced proportion of Dlxi1/2b::eGFP^+^ cells, we detected a strong reduction of *DLX2* expression in both *SMAD4* and *CSDE1* KO hSO at 35–40 days of differentiation (**Fig. 2f**, **P = 0.0032, ***P = 0.0005). In addition, *SMAD4* and *CSDE1* KO hSO showed a switch in ventral forebrain identity with reduced expression of several MGE-expressed transcription factors (*NXK2*.*1, LHX6, LHX8*, and *SOX6*^43,44^) and increased expression of caudal ganglionic eminence (CGE) and preoptic area (POA)-related transcription factors (*NR2F2, PROX1*, and *DBX1*^*43-45*^) (**Fig. 2f**, P < 0.03). Altogether, these results demonstrated that deletion of *SMAD4* and *CSDE1* reduces the proportion of Dlxi1/2::eGFP^+^ lineage cells in hSO, which likely results from disrupted subpallium patterning.

**Figure 2:**
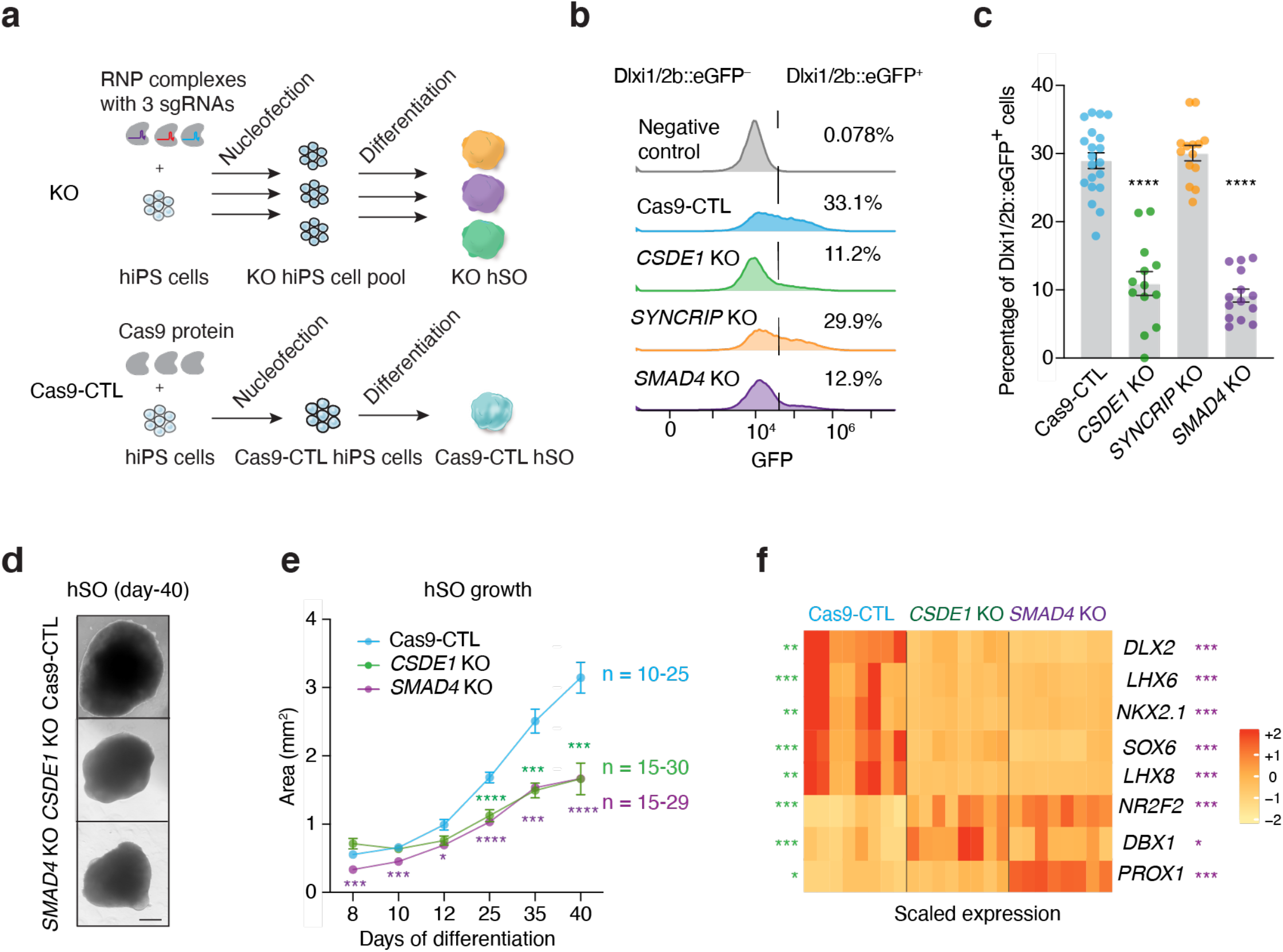
*CSDE1* and *SMAD4* regulate human interneuron generation. **a**. Schematic showing the generation of KO cell pools to validate candidate screen genes. Three sgRNAs per gene were used. Pre-mixed CAS9 protein and sgRNAs (RNP complex) were nucleofected into the CAG::Cas9;Dlxi1/2b::eGFP hiPS cells. Cells were cultured for several days, differentiated, and the dissociated at day 45 for flow cytometry analysis. Cells that were nucleofected with the Cas9 protein alone were used as CAS9-CTL. **b**, Histograms of flow cytometry analysis showing percentages of Dlxi1/2b::GFP^+^ cells in day 45 hSO. The negative control is control hSO without GFP expression. The X axis shows fluorescence intensity of GFP and the Y axis shows counts. **c**, Percentage of Dlxi1/2b::GFP^+^ cells in day 45 hSO from Cas9-CTL and KO samples (Cas9-CTL, n = 20, *CSDE1* KO, n = 13, *SYNCRIP1* KO, n = 14, and *SMAD4* KO, n = 14, from 2 independent nucleofections and differentiations). Each organoid was processed as one sample. Data are presented as mean ± SEM (standard error of mean). **** P < 0.0001; one-way ANOVA (F_3,57_ = 78.64) using Dunnett’s multiple comparison test. **d**, Bright field images of day 40 Cas9-CTL and KO hSO organoids. Scale bar: 400 μm. **e**, Area of hSO in the bright field images was measured for the Cas9-CTL and KO hSO from 8 to 40 days of differentiation (Cas9-CTL, n = 10–25, *CSDE1* KO, n = 15– 30, and *SMAD4* KO, n = 15–29, from 4 independent differentiations). Data are presented as mean ± SEM; mixed model two-way ANOVA (genotype as factor, F_2,81_ = 45.29) using Dunnett’s multiple comparison test. For the comparison of *CSDE1* KO and Cas9-CTL, P = 0.1647, 0.9191, 0.0559, <0.0001, 0.0001, 0.0002 at respective time points. For the comparison of *SMAD4* KO and Cas9-CTL, P = 0.0007, 0.0002, 0.0115, <0.0001, 0.0002, <0.0001 at respective time points. **f**, Heat map showing the differential expression of transcription factors in 35–40 days of differentiation Cas9-CTL and KO hSO examined by RT-qPCR (n = 8, from 4 independent differentiations); Mann Whitney test with Benjamini-Hochberg adjusted P-value. The stars on the left of the heatmap indicate the comparisons between Cas9-CTL and *CSDE1* KO. The stars on the right of the heatmap indicate the comparisons between Cas9-CTL and *SMAD4* KO. A detailed summary of adjusted P values for each analysis can be found in Supplementary Table 5.

### Loss of *TERF2* and *LNPK* impairs interneuron migration in forebrain assembloids

We next inspected the list of NDD gene hits for interneuron migration with a focus on genes that have not been generally associated with cell migration. Interestingly, we found *TERF2* a component of the shelterin complex that regulates telomere length^46^, and *LNPK*, which is proposed to serve as a curvature-stabilizing protein within tubular three-way junctions of the endoplasmic reticulum (ER)^47,48^ (**Fig. 1d** and **Supplementary Fig. 5a**). To verify whether these genes modulate interneuron migration, we engineered the CAG::Cas9;Dlxi1/2b::eGFP hiPS cells to generate KO cell pools for both *TERF2* and *LNPK* and confirmed the deletion of both genes (**Supplementary Fig. 5b–f**). Unlike hits from the first screen, loss of *TERF2* and *LNPK* did not affect the proportions of Dlxi1/2b::eGFP^+^ cells in hSO at 57 days of differentiation (**Supplementary Fig. 5g**), suggesting that they do not affect interneuron generation. We then measured fluorescence intensity of Dlxi1/2b::eGFP in unlabeled hCO fused with KO or Cas9-CTL hSO to estimate cellular migration (**Fig. 3a, Supplementary Video1**). Compared to Cas9-CTL hFA, we observed reduced mean intensity of GFP in the hCO assembled with the *TERF2* or *LNPK* KO hSO (**Fig. 3b, c**, Cas9-CTL vs *TERF2* KO: *P = 0.0225; Cas9-CTL vs *LNPK* KO: *P = 0.0428). This suggests that loss of *TERF2* or *LNPK* led to reduced migration of Dlxi1/2b::eGFP^+^ cells in forebrain assembloids.

**Figure 3:**
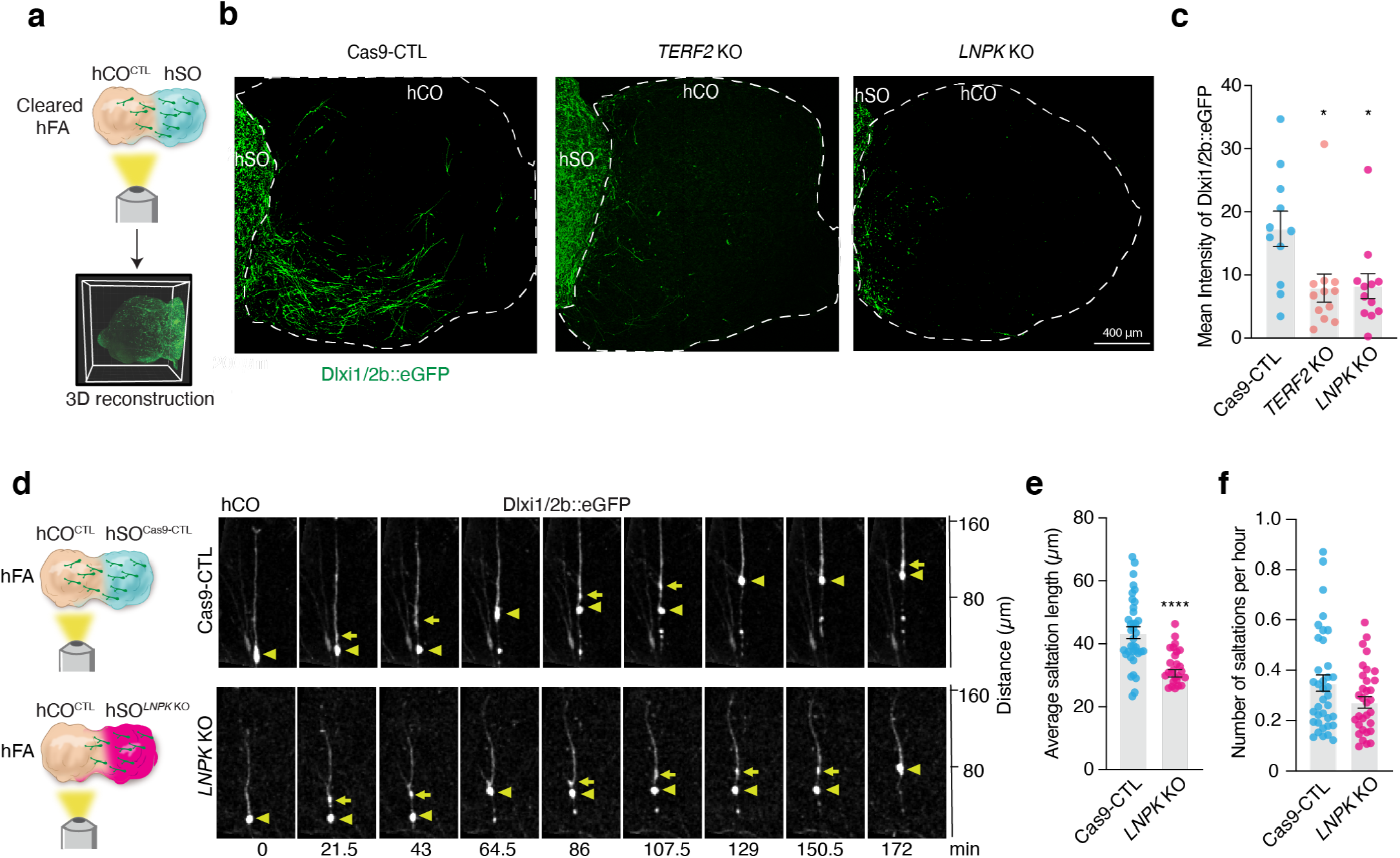
Deletion of *TERF2* and *LNPK* impairs interneuron migration in hFA. **a**. Schematic showing CUBIC-cleared hFA imaging and 3D reconstruction. **b**, Representative images of 2D projections from reconstructed 3D images of hFA (the entire hCO and part of hSO) showing hSO derived Dlxi1/2b::eGFP^+^ cells that have migrated into hCO. **c**, Mean intensity of GFP in the hCO side of the hFA (Cas9-CTL, n = 11 hFA; *TERF2* KO, n = 12 hFA; and *LNPK* KO, n = 12 hFA, from 2 independent differentiations). Data are presented as mean ± SEM. Cas9-CTL vs *TERF2* KO: *P = 0.0225; Cas9-CTL vs *LNPK* KO: *P = 0.0428, Kruskal-Wallis test with Dunn’s multiple comparisons test. **d**, Saltatory migration of Dlxi1/2b::eGFP^+^ cells in the hCO side of the Cas9-CTL and *LNPK* KO hFA. Triangles mark the nucleus and arrows mark the dilation that moves to the leading branch before nuclear translocation. **e–f**, Saltation length and number of saltations of Dlxi1/2b::eGFP^+^ cells in Cas9-CTL and *LNPK* KO hFA (n = 38 cells for each from 11 Cas9-CTL and 13 *LNPK* KO hFA from 4 independent differentiations). Data are presented as mean ± SEM; e, **** P < 0.0001, two-tailed unpaired t-test. f, P = 0.1332, Mann Whitney test.

### Abnormal migration of cortical interneurons following loss of the ER-related protein LNPK

Homozygous loss-of-function mutations in *LNPK* have been reported in patients with severe epilepsy^49^, which could involve abnormal development of cortical interneurons. To investigate the role of LNPK in interneuron migration into the cerebral cortex, we performed live imaging to examine in detail the movement of *LNPK* KO interneurons in hFA. Saltatory migration of interneurons involves, at the cellular level, two phases: first, the centrosome and the Golgi apparatus migrate forward in the leading branch and form a swelling; then, the nucleus translocates towards the swelling^50-52^. We monitored Dlxi1/2b::eGFP^+^ cells in the hCO side of hFA generated by fusing unlabeled hCO with Cas9-CTL or *LNPK* KO hSO, and we discovered reduced saltation length in *LNPK* KO migratory eGFP^+^ cells compared with that in Cas9-CTL (**Fig. 3d–f**, ****P < 0.0001, **Supplementary Video 2**). Taken together, this indicates that Dlxi1/2b::eGFP^+^ cells migrate less efficiently when LNPK is genetically removed.

While the role of the Golgi apparatus and the centrosome in interneuron migration has been extensively studied^50,52^, the ER has not been specifically described or directly implicated in this cellular process. This surprising ER-related gene hit (**Supplementary Fig. 5h**) prompted us to investigate ER dynamics in migrating human cortical interneurons. To visualize ER in migrating interneurons, we used a SEC61B-mEGFP hiPS cell line that labels the ER with mEGFP^53^. We performed confocal live imaging in hFA generated by fusing SEC61B-mEGFP hSO with unlabeled hCO to capture hSO-derived migratory cells. We found that a large fraction of the cellular ER is displaced along the leading branch in migrating cells. The ER takes on a linear conformation during this process before forming a compact structure in front of the nucleus, and this precedes nuclear translocation (**Fig. 4a, b, Supplementary Fig. 6** and **Supplementary Video 3**). We observed this phenomenon in 75% of saltatory migrating cells in the hSO side of the hFA and in 88% of cells in the hCO side (*P = 0.041; **Fig. 4c** and **Supplementary Fig. 7**). We further confirmed our findings in fixed hFA, which showed linear or compact ER in SEC61B-mEGFP^+^ cells that migrated to the hCO side of the hFA (**Supplementary Fig. 8a**). To confirm that this displacement is present in migrating interneurons, we virally labeled hFA comprised of SEC61B-mEGFP hSO and unlabeled hCO with Dlxi1/2b::mScarlet to label interneurons. We found that ER in Dlxi1/2b::mScarlet^+^ saltatory migrating cells showed a similar ER movement to the one described above (**Supplementary Fig. 8b, c**).

**Figure 4:**
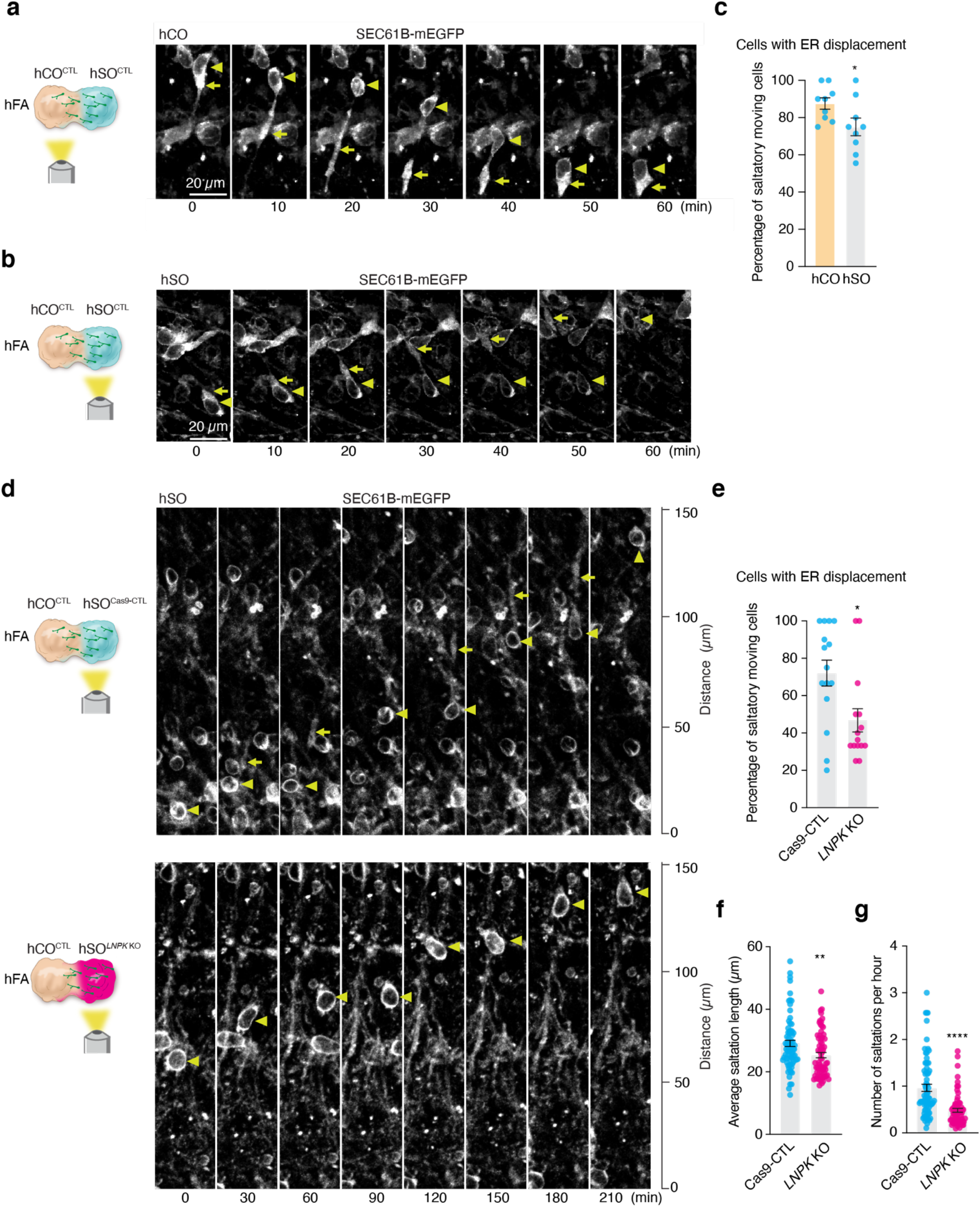
Deletion of LNPK impairs ER forward migration during nucleokinesis. **a–b**, Representative time-lapse sequences of SEC61B-mEGFP^+^ cells moving in a saltatory pattern in the hCO (**a**) and hSO (**b**) sides of the hFA (hCO is unlabeled). Images were taken at 59–80 days after hFA generation. Triangles mark the nucleus and arrows mark the linear or dilated structure of the ER during migration. The cell indicate in (**b**) also shows aggregation of ER signal at the rear of the soma. **c**, Percentages of saltatory moving cells imaged in the hCO or hSO sides of the hFA (at 59–80 days post-assembly) showing ER displacement (n = 9 hFA for each from 2 independent differentiations). Data are presented as mean ± SEM, *P = 0.041, Mann-Whitney test. **d**, Representative time-lapse sequences of SEC61B-mEGFP^+^ cells moving in a saltatory pattern in Cas9-CTL or *LNPK* KO hSO (in the SEC61B-mEGFP hiPS parental cell line), which was assembled with an unlabeled hCO. ER displacement was detectable in the last three steps of the four in Cas9-CTL, but not in the *LNPK* KO. **e**. Percentage of SEC61B-mEGFP^+^ saltatory moving cells in the hSO side of the hFA showing ER displacement (hFA at 29–52 days after assembly. n = 15 hFA from one differentiation). Data are presented as mean ± SEM. * P = 0.0218, Mann-Whitney test. **f–g**, Saltation length and number of saltations of SEC61B-mEGFP^+^ cells in (**e**) (Cas9-CTL, n = 73 cells; *LNPK* KO, n = 68 cells). Data are presented as mean ± SEM. ** P = 0.0042, **** P < 0.0001 Mann-Whitney test.

To investigate how loss of LNPK may affect ER displacement during interneuron migration, we generated *LNPK* KO hiPS cell pools (**Supplementary Fig. 9)** with the SEC61B-mEGFP hiPS cell line and then compared ER movement of SEC61B-mEGFP^+^ migrating cells in *LNPK* KO to Cas9-CTL hSO. *LNPK* KO hSO had a smaller proportion of cells displaying the forward ER movement prior to nuclear translocation (**Fig. 4 d, e**, *P = 0.0218, **Supplementary Video 4**). Consistent with our previous experiments in the CAG::Cas9;Dlxi1/2b::eGFP hiPS cell line, we observed reduced saltation length in *LNPK* KO SEC61B-mEGFP^+^ cells compared to Cas9-CTL (**Fig. 4f**, **P = 0.0042). In addition, we also found a reduction in the saltation frequency of the *LNPK* KO cells (**Fig. 4g**, ****P < 0.0001), which is likely related to the population of cells or the side of the assembloid that was imaged (versus **Fig. 3f**).

These data suggest that the ER migrates forward in the leading branch prior to nuclear translocation in migrating human cortical interneurons. Loss of LNPK results in a reduction in the proportion of cells undergoing ER displacement and may contribute to less efficient migration of *LNPK* KO interneurons in forebrain assembloids. Taken together, these results suggest that ER displacement prior to nucleokinesis is an essential step in human cortical interneuron migration.

## Discussion

Here, we have demonstrated a robust strategy to perform CRISPR loss-of-function screens for over 400 disease genes in hiPS cell-derived forebrain assembloids. This platform enabled us to systematically map loss-of-function phenotypes for NDD genes onto stages of human interneuron development. Notably, it highlighted the role of cytoskeleton machinery and, surprisingly, the ER in migration. Mutations in *LNPK*, which encodes an ER stabilizing protein^47,48^, cause severe intellectual disability (ID) and epilepsy in patients^49^, and we found that loss of LNPK in human interneurons resulted in defects in saltatory movement. This prompted us to further investigate the dynamics of the ER during migration, and we discovered a displacement of the ER in the leading branch that precedes nucleokinesis. ER-tubules interact with microtubules (MT)^54,55^, which are critical for establishing the leading process of migrating interneurons^56-59^. It could be that the disrupted ER-tubules structure following loss of LNPK^47,48^ interferes with the formation of cytoplasmic dilation in the leading neural process, which leads to a defect in cortical interneuron migration. More broadly, these experiments illustrate how mapping a list of disease-associated genes onto cellular pathways and specific stages of human brain development could ultimately identify convergent and divergent molecular and cellular phenotypes for these conditions and facilitate therapeutic efforts^60-62^.

There are several limitations to our study. First, the requirement of long-term cultures and assembling organoids impacts the number of genes that can be tested. Automation and strategies to accelerate interneuron maturation could facilitate the scaling and the cellular features that can be screened. Second, we used hSO that resemble the medial ganglionic eminences^21^, and considering the high diversity of GABAergic interneurons, there are likely cell-specific effects of NDD genes that have not been covered by these screens. Third, there are other non-neuronal cells that actively participate in interneuron generation and migration, such as vascular endothelial cells^63,64^ and oligodendrocytes^65^. The current platform does not capture non-cell autonomous effects on migration mediated by loss of NDD genes in these cells. Lastly, interneurons activate a program of maturation post-migration and integrate into circuits with glutamatergic neurons^17,66,67^. Future studies should screen effects on circuit assembly by employing assays that capture changes in activity following migration.

Building on the platform described here, systematic approaches to map a large group of disease genes onto human neural development and circuit function will be essential in revealing the biology of complex neurodevelopmental disorders and developing effective therapeutics.

## Methods

### Culture of hiPS cells

The hiPS cell lines used in this study were validated as previously described^68^. The SEC61B-mEGFP hiPS cell line (AICS-0059 cl.36) was developed at the Allen Institute for Cell Science (allencell.org/cell-catalog) and is distributed through Coriell^53^. The CAG::Cas9 cell line was genetically engineered in the lab (the parental cell line 1205-4 was derived and characterized at Stanford University^21^). The cell line was initially generated such that Cas9 expression can be turned off by adding doxycycline, but doxycycline was not used in this screen. The CAG::Cas9; Dlxi1/2b::eGFP cell line was generated in the CAG::Cas9 line. The 2242-1control hiPS cell line used to generate hCO was derived at Stanford University.

Briefly, hiPS cells were maintained in 6-well plates coated with recombinant human vitronectin (VTN-N, Life Technologies, A14700) in Essential 8 medium (Life technologies, A1517001). To passage hiPS cell colonies, cells were incubated with 0.5 mM EDTA for 7 minutes at room temperature, resuspended in Essential 8 medium and distributed into new 6-well plates. Cultures were tested and maintained Mycoplasma free. Approval for the derivation and used of these lines was obtained through Stanford IRB. Validation of hiPS cell genome integrity was performed by high density SNP arrays.

### Generating a list of NDD genes for the screen

We compiled a list of 611 genes associated with ASD and other NDDs based on genetic studies of NDD patients^8-11^ and the Simons Foundation Autism Research Initiative (SFARI) database (score 1, 2, and syndromic). To focus on NDDs genes that are expressed in hSO, we assessed gene expression in single-cell RNA-sequencing data from Dlxi1/2b::eGFP^+^ cells^21^. To ensure a conservative threshold, we averaged single-cell data from each of the collected samples (from assembloids of hCO and hSO) and plotted the maximum value across samples for each measured gene. We focused on the 425 NDD genes (out of 611 total) that showed at least minimal (defined as greater than 2 read counts) expression in at least one averaged sample. To confirm expression in the developing human forebrain, we also analyzed bulk RNA-sequencing data from the ganglionic eminences (MGE, CGE, and LGE) of developing primary human brain at 8-9 post-conception weeks (PCW) using the BrainSpan dataset^69^ (https://pubmed.ncbi.nlm.nih.gov/30545854/). Specifically, we obtained the maximum averaged expression value (log_2_ RPKM) across these samples, which is plotted in **Fig. 1a**. To visualize developmental expression trajectories of NDDs genes, normalized RPKM (obtained from http://development.psychencode.org/) expression values were log_2_ transformed and scaled (using the scale() function in R version 4.1.2 with default parameters). Data smoothing was performed using the geom_smooth() (as implemented in the ggplot2 package vergion 3.3.6) function in R with default parameters. Briefly, this function fits a penalized cubic regression spline to the data using the restricted maximum likelihood approach (via calling the mgcv:gam() R function with formula = y ∼ s(x, bs = “cs”) and method=“REML”).

### Generation of the CAG::Cas9 hiPS cell line

The CAG::Cas9 line was generated as previously described^70^. The donor plasmids were constructed by modifying the *AAVS1*-CAG-hrGFP (Addgene plasmids #52344) and replacing the GFP with Cas9 (Addgene plasmids #42230), TRE and tTA (Takara, Catalog No. 63113). The plasmid containing gRNA targeting *AAVSI* locus was obtained from Addgene (plasmid #41818). hiPS cells (2 × 10^6^ cells) dissociated with accutase (Innovative Cell Technologies, AT104) were electroporated using the P3 Primary Cell 4D-NucleofectorTMX Kit L (Lonza, V4XP-3024), a 4D-nucleofector core unit and X unit (Lonza; nucleofection program CA-137). Cells were seeded on a well of a 6-well plate that was pre-coated with vitronectin and contained pre-warmed Essential 8 medium supplemented with the ROCK inhibitor Y-27632 (10 μM; Selleckchem, S1049). Once the seeded cells reached 70–80% confluency, 1 μg/ml of puromycin in StemFlex medium (Life Technologies, A3349401) was supplemented for 5 days. Colonies were visible at the end of 5 days of selection in puromycin. Cells were maintained in StemFlex medium for another 5 days, and individual colonies were picked and maintained separately in a 24-well plate coated with vitronectin. Cells were maintained in StemFlex medium for another 4 days and were expanded and maintained in Essential 8 medium afterwards. Three sets of PCR programs were designed to select colonies that showed the insertion of the left, middle, and right parts of the donor DNA (PCR1: 5’-TCCTGAGTCCGGACCACTTT and 3’-CACCGCATGTTAGAAGACTTCC; PCR2: 5’-TATGGAGATCCCTCGACCTG and 3’-CCTGGGATACCCCGAAGAGT; PCR3: 5’-TTCTTCTTGATGCTGTGCCG and 3’-GCTCAAGGGGCTTCATGATG). Validation of hiPS genome integrity was performed by high density SNP arrays.

### Generation of the CAG::Cas9;Dlxi1/2b::eGFP cell line

CAG::Cas9 hiPS cells cultured in a 6-well plate were infected with lentivirus expressing Dlxi1/2b::eGFP with a hygromycin resistance gene. Cells were maintained in Essential 8 medium for another 3 days before passaging. One day after passaging, 200 μg/ml hygromycin was added to the medium to select for infected cells. Cells were maintained in hygromycin for 9 days. Control cells without virus infection died after 4 days of exposure to 200 μg/ml hygromycin. Surviving cells were expanded and cryopreserved for use.

### Generation of the interneuron migration CRISPR sgRNA library

The interneuron migration CRISPR sgRNA library was designed by selecting up to 5 sgRNAs targeting each of the 438 genes of interest (425 disease genes and 13 genes associated with interneuron development) and an additional 10% ‘safe’ harbor-targeting negative control sgRNAs (Supplementary Table 2)^71^. Cloning restriction sites and amplification primer handles were appended to each oligo. The sgRNA library oligos were synthesized on the Agilent chip oligo array synthesis platform and cloned into a lentiviral plasmid vector that constitutively expresses the sgRNA and mCherry.

### Generation of hCO and hSO from hiPS cells

hCO were differentiated from feeder-free maintained hiPS cells, as previously described^68^. To generate 3D hCO, hiPS cells were dissociated with accutase to obtain a single-cell suspension. About 3 × 10^6^ cells were seeded per AggreWell™800 plate well in Essential 8 medium supplemented with the ROCK inhibitor Y27632 (10 μM). The plate was then centrifuged at 100 x g for 3 minutes and incubated overnight at 37°C with 5% CO_2_. The next day (day 0), cellular aggregates were transferred into ultra-low attachment plastic dishes (Corning, 3262) and cultured in Essential 6 medium (Thermo Fisher Scientific, A1516401) supplemented with dorsomorphin (2.5 μM, Sigma-Aldrich, P5499), SB-431542 (10 μM, R&D Systems, 1614), and XAV-939 (1.25 μM, Tocris, 3748). No medium change was performed on day 1. Organoids were cultured in E6 medium with patterning molecules from day 2–5. On day 6, organoids were transferred to neural medium containing Neurobasal A (Thermo Fisher, 10888022), B-27 supplement without vitamin A (Life Technologies, 12587), GlutaMax (1:100, Life Technologies, 35050061), and penicillin-streptomycin (1:100, Thermo Fisher, 15140163). Neural medium was supplemented with 20 ng/ml EGF (R&D Systems) and 20 ng/mL FGF2 (R&D Systems) from day 6 to day 24 with daily medium change in the first 10 days, and every other day for the subsequent 9 days. From day 25 to day 43, neural medium was supplemented with 20 ng/ml BDNF (Peprotech, 450-02) and 20 ng/ml NT-3 (Peprotech, 450-03). From day 44, only neural medium without growth factors was used for medium changes every 4 days. Gellan gum (∼0.005%, Sigma, G1919-259G, lot SLCF5726) was added in the medium from day 6–15 to avoid spontaneous organoid fusion^72^. The differentiation of hSO from hiPS cells is similar to hCO differentiation with the following differences in patterning: from day 6 to day 11, EGF, FGF2, and XAV-939 (1.25 μM, Tocris, 3748) were supplemented to the neural medium; from day 12 to day 24, EGF, FGF2, XAV-939, and SAG (100nM, EMD Millipore, 566660) were used. Gellan gum (∼0.005%) was added in the medium from day 6–11 to avoid spontaneous organoid fusion.

### Generation of hFA

To generate hFA, hCO and hSO were generated separately and assembled by placing them in close proximity in 24-well low attachment plates (Corning 3473), which were placed slightly tilted in the incubator^73^. Medium was changed 3 days later. Assembloids were then cultured in the 24-well low attachment plates until use, with the plates placed flat in the incubator. For the screen and the validations, day 45 hSO were fused with day 60 hCO, and hFAs were used for downstream analysis at ∼30 days post-assembly. For the experiments imaging ER morphology, day–60 hSO and hCO were assembled as previously described^21^ and hFA were imaged at specific days post-assembly as described.

### Interneuron generation and migration screen

CAG::Cas9;Dlxi1/2::eGFP hiPS cells were passaged with accutase one day before infection with the lentiviral library (with 5 μg/ml polybrene). Three days later, mCherry^+^ cells were selected by FACS (collected ∼3.6 × 10^6^ cells) and seeded onto vitronectin coated plates to recover and expand for another 6 days. hSO were generated with these hiPS cells as per described above. At day 44, hSO were collected for the interneuron generation screen. At a similar time, hFA were generated by assembling day 60 hCO generated from an unlabeled hiPS cell line with day 45 hSO. We generated a total of 1008 assembloids. Thirty days post-assembly, hFA were checked for migration under a fluorescence microscope. hFA that didn’t pass quality control (i.e., displayed below average GFP signal on the hCO side) were excluded. hFA were then aligned on 100-mm plates with hSO on the left and hCO on the right, and were split at the boundary of the assembloid under a dissection scope. hCO and hSO were collected separately after checking GFP expression under a fluorescence microscope. The hSO showed strong GFP signal in the entire organoid while the hCO showed less GFP signal. The samples were subsequently dissociated, and populations were sorted out and stored in a – 80°C freezer. We processed ∼900 assembloids in 8 days. In addition, 5 × 10^6^ hiPS cells and day 19 organoids sampled from all plates were collected.

### Lentivirus production

Lentivirus was produced by transfecting HEK293T cells with lentiviral transfer plasmids, packaging plasmids, and envelope plasmids using PEI (Polysciences, 24765-100). Media from transfected cells was harvested at 24, 48 and 72 hours after transfection. Viruses were concentrated by spinning at 17,000 rpm at 6°C for 1 hour.

### Preparation of sequencing libraries

Sample-specific sequencing libraries were prepared from the extracted genomic DNA of each harvested sample by amplifying and barcoding the lentivirally-integrated sgRNA spacer sequence using the Herculase II Fusion DNA Polymerase (Agilent Cat. 600679). Samples were pooled and sequenced on Illumina NextSeq 550 flow cells at 450–1000X read depth.

### Analyzing screen results

Screen results were analyzed using the CasTLE software^71^– a maximum likelihood estimator that provides for each gene: a CasTLE score, a log-likelihood ratio that considers the relative enrichment or depletion of each gene-targeting sgRNA compared to the distribution of negative control sgRNA effects, a CasTLE effect size, a log_2_-transformed maximum likelihood estimate of the ratio of cells containing sgRNAs perturbing a given gene in one sample compared to another, normalized to the median difference of negative control sgRNAs. In the interneuron generation screen, genes with at least 2 sgRNAs having effects ≤ –1.5 or ≥ +1.5, approximately 2x the SD of negative controls, and no sgRNAs with opposite effects beyond the threshold, were selected as candidates. In the interneuron migration screen, genes with at least 2 sgRNAs having effects ≤ –2.5 or ≥ +2.5, approximately 2x the SD of negative controls, and no sgRNAs with opposite effects beyond the threshold, were selected as candidates.

### Generating KO cell pools with 3x sgRNAs strategy

To validate the candidate genes of the screen, we generated KO hiPS cell pools for genes of interest with the CRISPR/Cas9 system. Three sgRNAs targeting an early exon of a specific gene were designed and synthesized by Synthego (see **Supplementary Table 6** for the sgRNAs sequences) to induce one or more fragment deletions (**Supplementary Fig. 4b**). In brief, CAG::Cas9;Dlxi1/2::eGFP hiPS cells were dissociated with accutase and 0.5 million cells were mixed with 300 pmol sgRNAs and 40 pmol Cas9 protein (Synthego, SpCas9 2NLS Nuclease (1000 pmol)). Nucleofection was performed using the P3 Primary Cell 4D-NucleofectorTMX Kit L (Lonza, V4XP-3032), a 4D-nucleofector core unit and the X unit (Lonza) (program CA-137). Cells were then seeded onto vitronectin coated 6-well plates in Essential 8 medium supplemented with the ROCK inhibitor Y27632 (10 μM). Essential 8 medium was used for daily medium change.

### Preparing samples for next generation sequencing (NGS)

DNA was isolated from hSO using the DNeasy Blood & Tissue Kit (Qiagen, 69506). Primers (Supplementary Table 7) with Illumina adaptors were used to amplify ∼300 bp of DNA sequence around the sgRNA targeting site of the gene of interest. PCR was performed with the PrimeSTAR Max DNA polymerase 2x hot-start PCR master mix (TaKaRa Bio, R045A). PCR products were purified using the AMPure XP PCR purification system (Beckman Coulter, A63881) and the 96S Super Magnet (Alpaqua, A001322) for next generation sequencing (NGS, Azenta).

### Dissociation of hCO and hSO

hCO and hSO were dissociated as previously described^74^ with some modifications. Briefly, organoids (single organoids cultured in a 96-well plate or up to 8 organoids cultured in a 6-well plate) were incubated with an enzyme solution containing 10U/ml papain (Worthington, LS003127), 1x EBSS (Sigma-Aldrich, E7510-500ML), 0.36% D(+)-Glucose (Sigma-Aldrich), 26 mM NaHCO_3_ (Sigma-Aldrich), 0.5 mM EDTA (Sigma-Aldrich), DNase (Worthington, LS002007), 6.1 mM L-Cysteine (Sigma-Aldrich), and the ROCK inhibitor Y-27632 (10 μM) at 37°C with 5% CO_2_ for 15 minutes; this solution was pre-warmed at 37°C for 30 minutes to activate papain, and then cells were washed with a pre-warmed protease inhibitor solution (1x EBSS, 0.36% D(+)-Glucose, 26 mM NaHCO_3_, 0.2% trypsin inhibitor (Worthington LS003086)). Organoids were then triturated, and the resulting single cell suspension was centrifuged at 1,200 rpm for 5 minutes. The pellet was resuspended with 3% BSA (Sigma-Aldrich) and then filtered.

### Flow Cytometry

To sort mCherry^+^ or GFP^+^ cells during the screen, FACS was conducted on FACS-Aria II instruments (Stanford FACS Facility). The ACEA NovoCyte Quanteon 4025 flow cytometer (Stanford FACS Facility) was used to analyze the percentage of GFP^+^ cells. Data were analyzed on FlowJo (version 10.8.1).

### Real-time quantitative PCR (RT-qPCR)

For RT-qPCR analysis, 2–3 organoids were pooled as a sample. mRNA was isolated using the RNeasy Mini Plus kit (Qiagen, 74136) and template cDNA was prepared by reverse transcription (RT) using the SuperScript III First-Strand Synthesis SuperMix for qRT-PCR (Thermo Fisher Scientific, 11752250). RT-qPCR was performed using the SYBR Green PCR Master Mix (Thermo Fisher Scientific, 4312704) on a QuantStudio 6 Flex Real-Time PCR System (Thermo Fisher Scientific, 4485689). Data was processed using the QuantStudio RT-PCR software (Applied Biosystems). To plot the heatmap to present the qPCR results, gene expression was scaled and mean centered (using the “scale” function in R) and the heatmap was plotted using the “ComplexHeatmap” package. Primers used in this study are listed in Supplementary Table 8.

### Clearing, staining and imaging of hFA

The Clear, Unobstructed Brain/Body Imaging Cocktails and Computational analysis (CUBIC) method was applied to optically clear hFA, as previously described^75^. Briefly, hFA at ∼40 days post-assembly were fixed overnight with 4% PFA at 4°C. The next day, samples were washed with PBS (2 hours x 3, 37°C) before incubation with the Tissue-Clearing Reagent CUBIC-L (TCI, T3740) at 37°C for ∼18 hours. After washing with PBS (2 hours x 3, 37°C), samples were incubated in a nuclear staining solution (1:150 dilution of RedDot, Biotium 40061-T) in PBS with 500 mM NaCl) at 37°C overnight. Samples were then washed with PBS (2 hours x 2, 37°C) followed by incubation with HEPES-TSB (10 mM HEPES, 10% TX-100, 200 mM NaCl, 0.5% BSA) buffer at 37°C for 2 hours. Samples were subsequently stained with anti-GFP antibody (GTX13970, 1:1,000) for 48 hours and secondary antibody (Alexa Fluor® 488 AffiniPure Donkey Anti-Chicken IgY, Jackson ImmunoResearch 703-545-155, 1:300) for another 48 hours. Antibodies were prepared in HEPES-TSB buffer and hFA were washed with 10% TX-100 in PBS (2 hours x 2, 37°C) and HEPES-TSB (2 hours x 1, 37°C) in between stains. After secondary antibody staining, samples were washed with 10% TX-100 in PBS (30 minutes x 2, 37°C) and PBS (1 hour x 1, 37°C) followed by subsequent incubation with 50% and 100% Tissue-Clearing Reagent CUBIC-R+ (TCI, T3741) at room temperature for 1 day each. All incubation steps were performed on a shaker. Samples were transferred to a 96-well plate with a clear bottom (ibidi, 89626) in CUBIC-R+ solution and imaged using a 10x objective on a Leica Stellaris confocal microscope. The entire hCO and the hCO-hSO boundary were imaged (∼1 mm volume per hFA) blindly to the genotype of the samples.

Images were analyzed with Imaris (Oxford Instruments, v9.7.0) and Fiji (ImageJ, v1.0 and v1.53f51). Briefly, Z stack confocal images were compiled in Imaris to generate a 3D reconstruction of the hFA. The boundary of the assembloid in individual frames was manually delineated using the nuclear stain signal (every ∼40 frames); these boundaries were used to create an Imaris “surface” to define the hCO (Supplementary Video 1). The 3D hCO was then reduced into a 2D projection using the maximum intensity function in Fiji. To measure the Dlxi1/2b::eGFP intensity in a consistent manner for all samples, the hCO-hSO boundary was aligned vertically. One third of the hCO adjacent to the boundary was used to quantify mean intensity. A representative area of background noise was also measured for mean intensity and was subtracted from the measurements of the hCO.

### Western Blotting

Day 60–80 hSO were used for western blotting. Whole cell protein lysates were prepared using the RIPA buffer system (Santa Cruz, sc-24948); protein concentration was quantified using the Bicinchoninic Acid (BCA) assay (Pierce, ThermoFisher 23225). For electrophoresis, 20 μg of protein per lane was loaded and run on a 4–12% Bis-Tris PAGE gel (Bolt 4–12% Bis-Tris Protein Gel, Invitrogen, NW04122BOX) and transferred onto a PVDF membrane (Immobilon-FL, EMD Millipore). Membranes were blocked with 5% BSA in TBST for 1 hour at room temperature (RT) and incubated with primary antibodies against GAPDH (mouse, 1:5000, GeneTex, GTX627408) overnight and LNPK (rabbit, 1:500, Sigma, HPA014205) for 72 hours at 4°C. Membranes were washed 3 times with TBST and then incubated with near-infrared fluorophore-conjugated species-specific secondary antibodies: Goat Anti-Mouse IgG Polyclonal Antibody (IRDye 680RD, 1:10,000, LI-COR Biosciences, 926–68070) or Goat Anti-Rabbit IgG Polyclonal Antibody (IRDye 800CW, 1:10,000, LI-COR Biosciences, 926–32211) for 1 hour at RT. Following secondary antibody application, membranes were washed 3 times with TBST, once with TBS, and then imaged using a LI-COR Odyssey CLx imaging system (LI-COR).

### Immunofluorescence staining

hFA were fixed in 4% PFA-PBS overnight at 4°C followed by dehydration in 30% sucrose for 3 days. Subsequently, samples were embedded in optimal cutting temperature (OCT) compound (Tissue-Tek OCT Compound 4583, Sakura Finetek) and 30% sucrose–PBS (1:1) for cryosectioning (25 μm-thick sections) using a Leica Cryostat (Leica, CM1860). For immunofluorescence staining, cryosections were washed with PBS to remove excess OCT and blocked in 10% normal donkey serum (NDS, MilliporeSigma, S30-M) and 0.3% Triton X-100 (MilliporeSigma, T9284-100ML) diluted in PBS for 1 hour at room temperature. Sections were then incubated overnight at 4°C with primary antibodies diluted in PBS containing 10% NDS and 0.3% Triton X-100. PBS was used to wash away excess primary antibodies, and the cryosections were incubated with secondary antibodies in PBS containing 10% NDS and 0.3% Triton X-100 for 1 hour. The following primary antibodies were used for staining: anti-GFP antibody (1:1,000 dilution, chicken, GeneTex, GTX13970) and anti-LNPK antibody (1:500, rabbit, Sigma, HPA014205). Alexa Fluor 488 AffiniPure Donkey Anti-Chicken IgY (Jackson ImmunoResearch, 703-545-155) and Alexa Fluor 568 donkey anti-rabbit IgG (H&L) highly cross-adsorbed secondary antibody (Thermo Fisher Scientific, A10042) were used in 1:1,000 dilution. The nuclei were visualized with Hoechst33258 (Thermo Fisher Science, H3569, 1:10,000 dilution). Cryosections were mounted for microscopy on glass slides using Aquamount (Polysciences, 18606) and imaged on a Leica Stellaris confocal microscope. Images were processed with Fiji (ImageJ, v1.0)

### Live cell imaging and analysis of Dlxi1/2b::eGFP^+^ cell migration

To image the Dlxi1/2b::eGFP^+^ cell migration, hFA were transferred to a well of a 96-well plate with a clear bottom (ibidi, 89626) in 500 μl neural medium under environmentally controlled conditions (37°C, 5% CO_2_). Samples were placed in this condition for 30–60 minutes before imaging on a confocal microscope (Leica SP8). hFA were imaged under a 10x objective at a depth of 50–150 μm and at a rate of ∼20 minutes per frame for 18–22 hours. Fiji (ImageJ, v1.0) was used to quantify the migration of Dlxi1/2b::eGFP^+^ cells. To estimate the length of individual saltations, Dlxi1/2b::eGFP^+^ cells displaying a swelling of the soma were identified, and the distance (in μm) to the new position of the soma following nucleokinesis was measured manually. The time necessary for this movement was used to calculate saltation frequency. Only cells that showed at least 2 complete jumps during the imaging session were included in the analysis, and the averaged saltation length of each cell was used in the statistical analysis. Analysis was performed blindly to the genotype.

### Live cell imaging and analysis of SEC61B-mEGFP^+^ cell migration

To image the SEC61B-mEGFP^+^ cells, hFA (SEC61B-mEGFP^+^ CTL, CAS9-CTL, or *LNPK* KO hSO assembled with unlabeled hCO) were imaged under environmentally controlled conditions (37°C, 5% CO_2_) on a confocal microscope (Leica Stellaris, 20x objective, 2x zoom). The samples were placed in the controlled environment for 1 hour before imaging. Samples were imaged at a depth of ∼12 μm and a rate of 10 minutes per frame for ∼20 hours. Given the short working distance of the 20x objective and the nature of 3D floating samples, only images at the very bottom of the samples were used. One to three images per hFA were obtained for the analysis. Images that contain densely packed fast-moving cells were excluded. To analyze the percentage of saltatory migrating cells in hCO or hSO showing ER displacement, SEC61B-mEGFP^+^ cells displaying saltatory migrations with at least 2 jumps in the imaging session were selected, and cells displaying a swelling of the ER at the leading neural branch preceding nuclear translocation were identified as exhibiting ER displacement (cells were considered to have this ER displacement if one of the jumps showed this displacement). The hCO or hSO with at least three saltatory moving cells were included from the analysis. When estimating the saltation length and saltation frequency of SEC61B-mEGFP^+^ migratory cells (Fig. 4f, g), a strategy similar to that used for measuring Dlxi1/2b::eGFP^+^ migrating cells was employed. All cells used to generate Figure 4e with at least 2 complete jumps were included. Fiji (ImageJ, v1.0 and v1.53f51, and ImageJ2, v2.3.0/1.53q) was used to analyze the migration of SEC61B-EGFP^+^ cells. Blinding for genotype was used for analysis.

When examining Dlxi1/2b::mScarlet virally labeled hFA (SEC61B-mEGFP hSO assembled with unlabeled hCO), Dlxi1/2b::mScarlet^+^ cells were identified by the florescence signal filling the entire soma and extending to the leading branch (SEC61B-mEGFP hiPS line also expresses mTagRFP-T tagged LMNB1, which was captured while imaging mScarlet). ER displacement was examined based on SEC61B-mEGFP signal as described above.

### Viral labeling for live imaging

hFA (SEC61B-mEGFP hSO assembled with unlabeled hCO) were infected with lentivirus encoding Dlxi1/2::mScarlet. Briefly, 30 days post-assembly, hFA cultured in 24-well low attachment plates with 500 μl neural medium were cultured with virus overnight at 37°C with 5% CO_2_. The next day, fresh neural medium (500 μl) was added. The following day, the medium was replaced with fresh medium. Samples were ready for live imaging 2 weeks after infection.

## Statistics

Data are presented as mean ± SEM, unless otherwise indicated. Distribution of the raw data was tested for normality of distribution; statistical analyses were performed using the Student’s t-test, Mann Whitney test, Kruskal-Wallis test, and one-way or two-way ANOVA with multiple comparison tests as indicated. Sample sizes were estimated empirically. Blinding of genotypes was used for imaging analyses.

## Data Availability

The codes for casTLE analysis can be downloaded at: https://bitbucket.org/dmorgens/castle/downloads/.

All data and other codes are available from the corresponding author upon request.

## Author Contributions

X.M and S.P.P conceived the project and designed the experiments. X.M. conducted the differentiation, sample collection, and library preparation for the screen. K.K. analyzed the NDD genes and generated the list of genes used in the screen. D.Y designed and cloned the sgRNA library, sequenced the library and analyzed the data with the supervision of M.C.B. S.K. generated the lentivirus encoding the sgRNA library. X.C. performed FACS for the screen. X.M. and N.R. generated the KO cell pools, and performed flow cytometry, live imaging, and cell migration analysis. M.V.T. performed western blot. X.M. and S.P.P. wrote the manuscript with inputs from all authors. S.P.P. supervised all aspects of this study.

## Acknowledgements

We thank J. Bernstein, R. Reimer and members of the Paşca laboratory for scientific input, especially S. Kanton. This work was supported by the Stanford Brain Organogenesis Program in the Wu Tsai Neuroscience Institute and Bio-X, Kwan Funds, Senkut Funds, the Ludwig Foundation, the Mann Foundation (SPP), and the Stanford Medicine Dean’s Fellowship (to XM). SPP is a New York Stem Cell Foundation (NYSCF) Robertson Stem Cell Investigator, a Chan Zuckerberg Initiative (CZI) Ben Barres Investigator and a CZ BioHub Investigator. Flow cytometry analysis and cell sorting were performed on instruments in the Stanford Shared FACS Facility obtained using NIH S10 Shared Instrument Grant (S10RR025518-01). This paper was typeset with the bioRxiv word template by @Chrelli: www.github.com/chrelli/bioRxiv-word-template.

## Competing Interests

Stanford University holds multiple patents on organoids and assembloids with S.P.P. listed as an inventor.

**Supplementary Figure 1:**
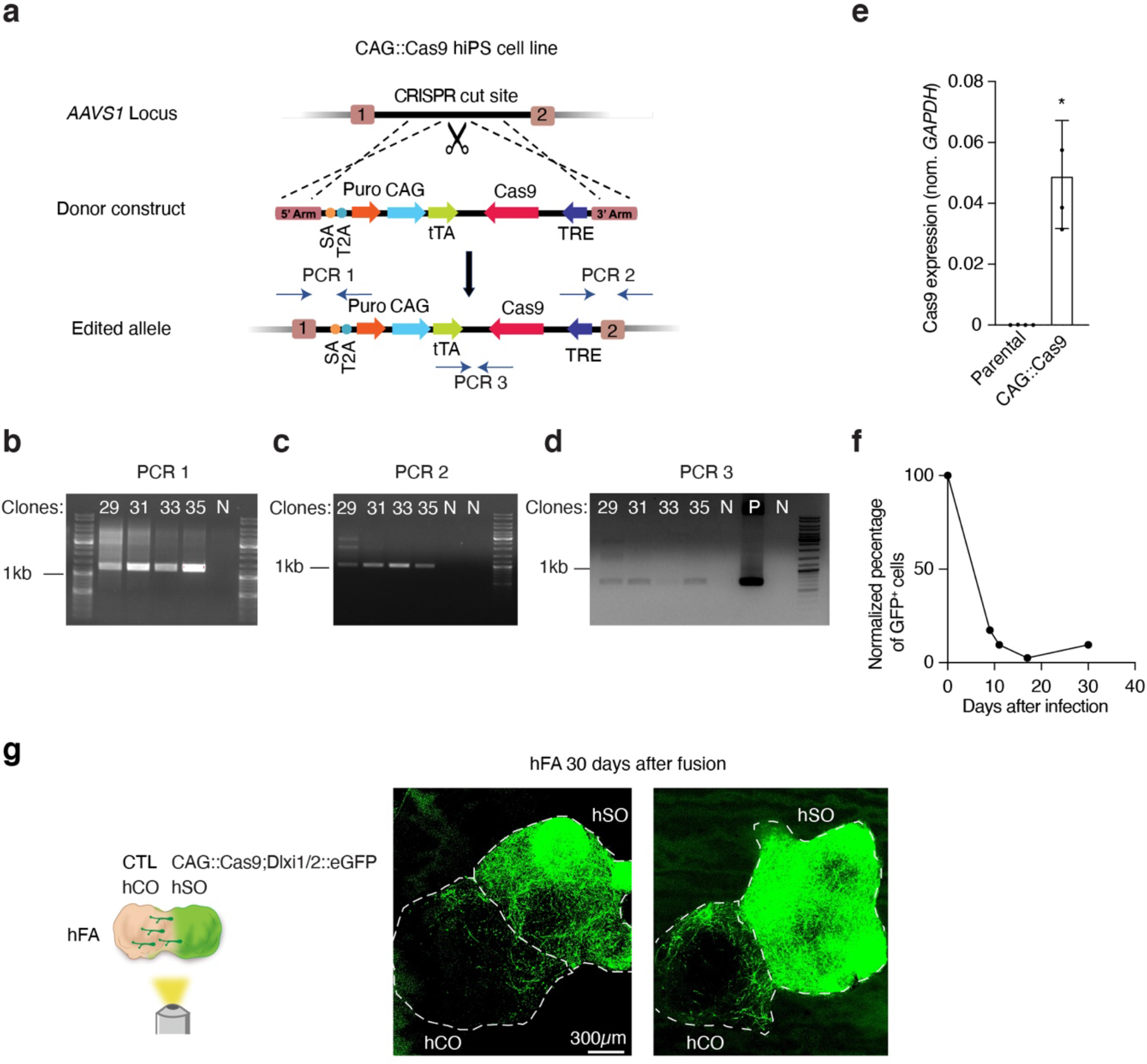
Generation and validation of the CAG::Cas9 hiPS cell line. **a**. Schematic showing the strategy to generate the CAG::Cas9 hiPS cell line. Dashed lines indicate homologous recombination sites. PCR 1–3 indicate the three pairs of primers designed to confirm the insertion of the entire donor. **b–d**. PCR confirming the successful amplification of the targeted DNA as shown in (**a**) to identify the clone with complete insertion of the donor construct. “N” indicates a negative control, which is the PCR product with genomic DNA extracted from a control hiPS cell line. “P” indicates a positive control, which is the PCR product from the donor plasmid. In total, 35 clones were screened, and 3 clones (Clone #29, #31, and #35) were positive in all PCR reactions. **e**. Cas9 expression in CAG::Cas9 hiPS cells compared to the parental cells (n = 4) by RT-qPCR. Data are presented as mean ± SEM; * P = 0.026, Mann Whitney test. **f**. Genome editing efficiency in CAG::Cas9 hiPS cells. CAG::Cas9 hiPS cells were infected with a lentivirus harboring GFP and a sgRNA against GFP. Infected cells were split into two groups with or without doxycycline (DOX) supplementation to inhibit Cas9 expression. Cells were subjected to flow cytometry to examine the percentage of GFP^+^ cells. The percentage of GFP^+^ cells (–DOX)/percentage of GFP^+^ cells (+DOX) at each time point was plotted. **g**. Representative live images of hFA at 30 days after assembly. Day 45 CAG::Cas9;Dlxi1/2b::eGFP hSO were assembled with day 60 unlabeled hCO. The dashed lines indicate the boundary of hCO and hSO.

**Supplementary Figure 2:**
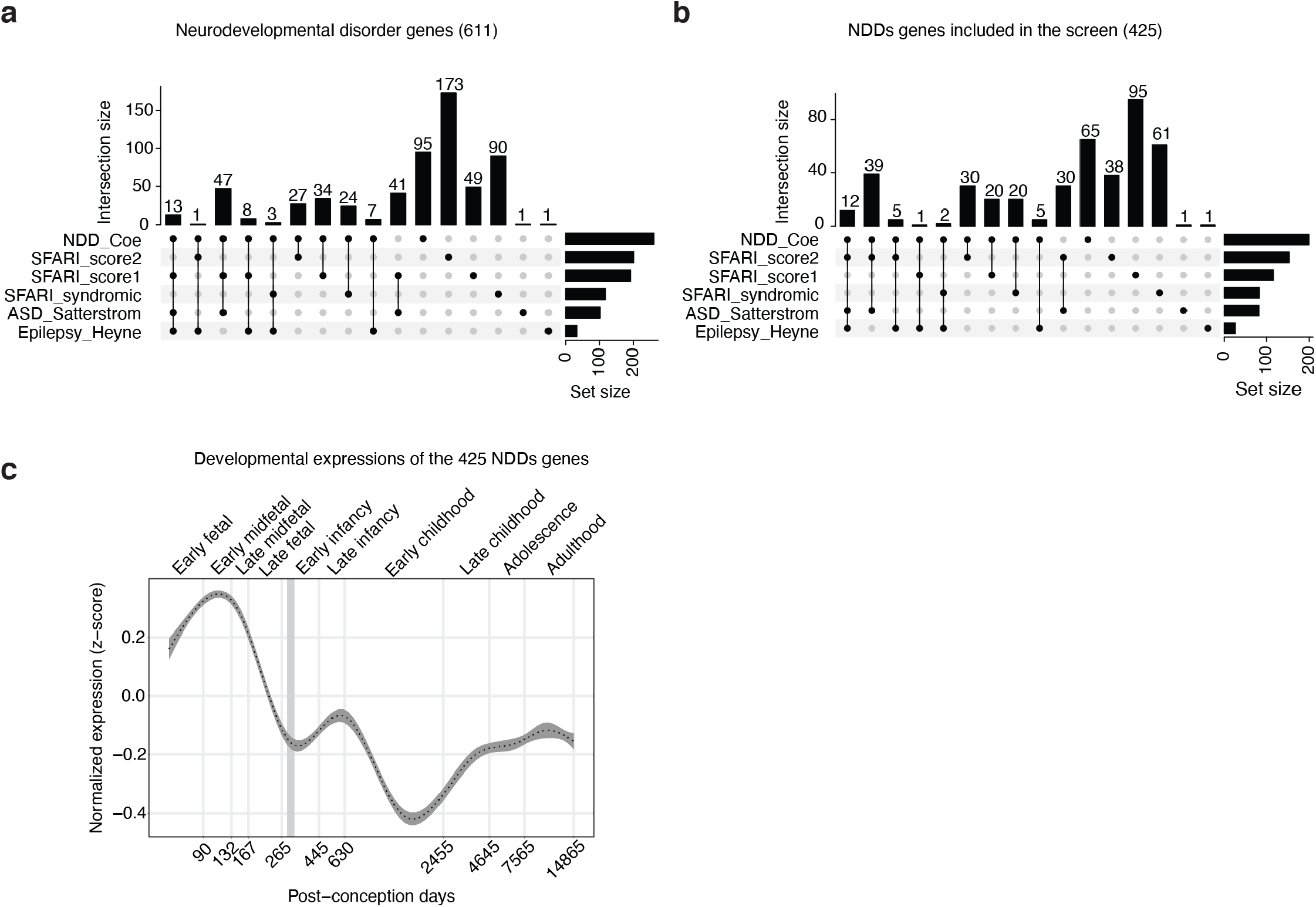
Summary of NDD genes and the genes included in the screen. **a**. Source of the 611 NDD genes. **b**. Source of the 425 NDD genes included in the screens. **c**. Expression trajectories in the human cerebral cortex of the selected 425 NDD genes using the BrainSpan dataset^69^. Displayed are smoothened (generalized additive model) z-score normalized expression data with 99% confidence interval shown in gray (n = 425 genes; n = 419 cortical samples).

**Supplementary Figure 3:**
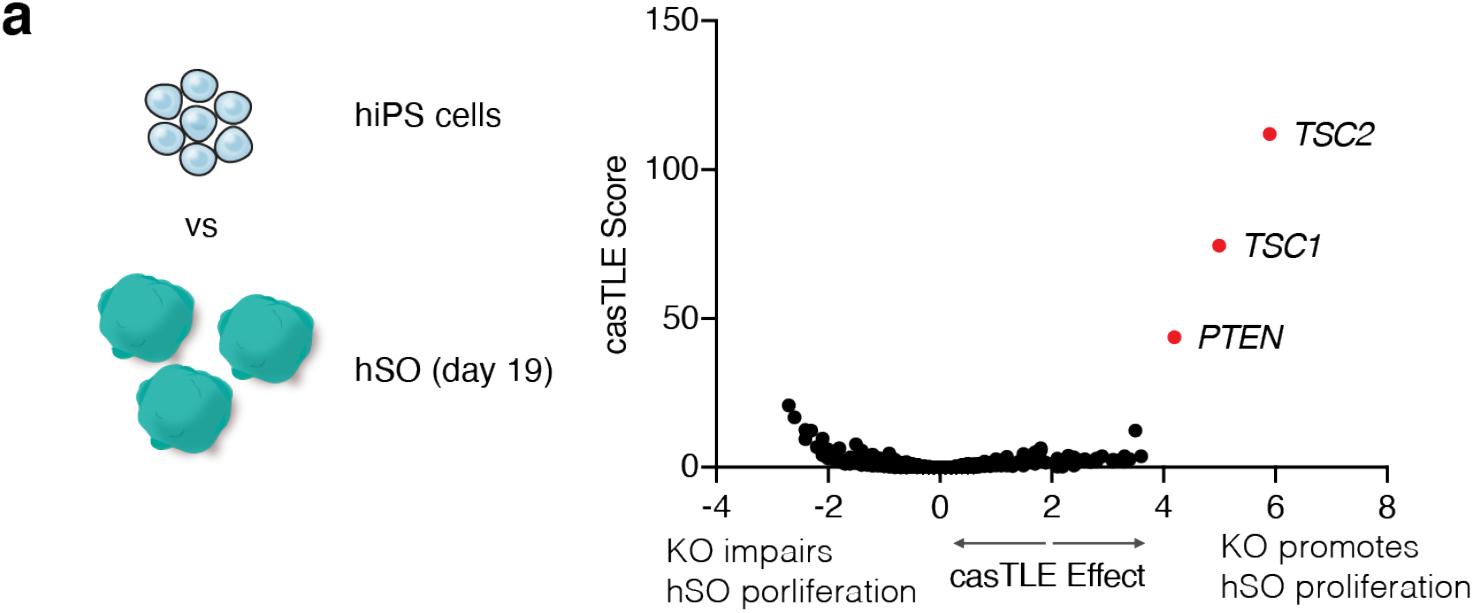
Early neural differentiation genes captured in the screen. **a**. Volcano plot of the CasTLE-estimated maximum gene perturbation effect size and associated CasTLE score for neural differentiation at 19 days of hSO differentiation. Three genes with notable effects are labeled in red.

**Supplementary Figure 4:**
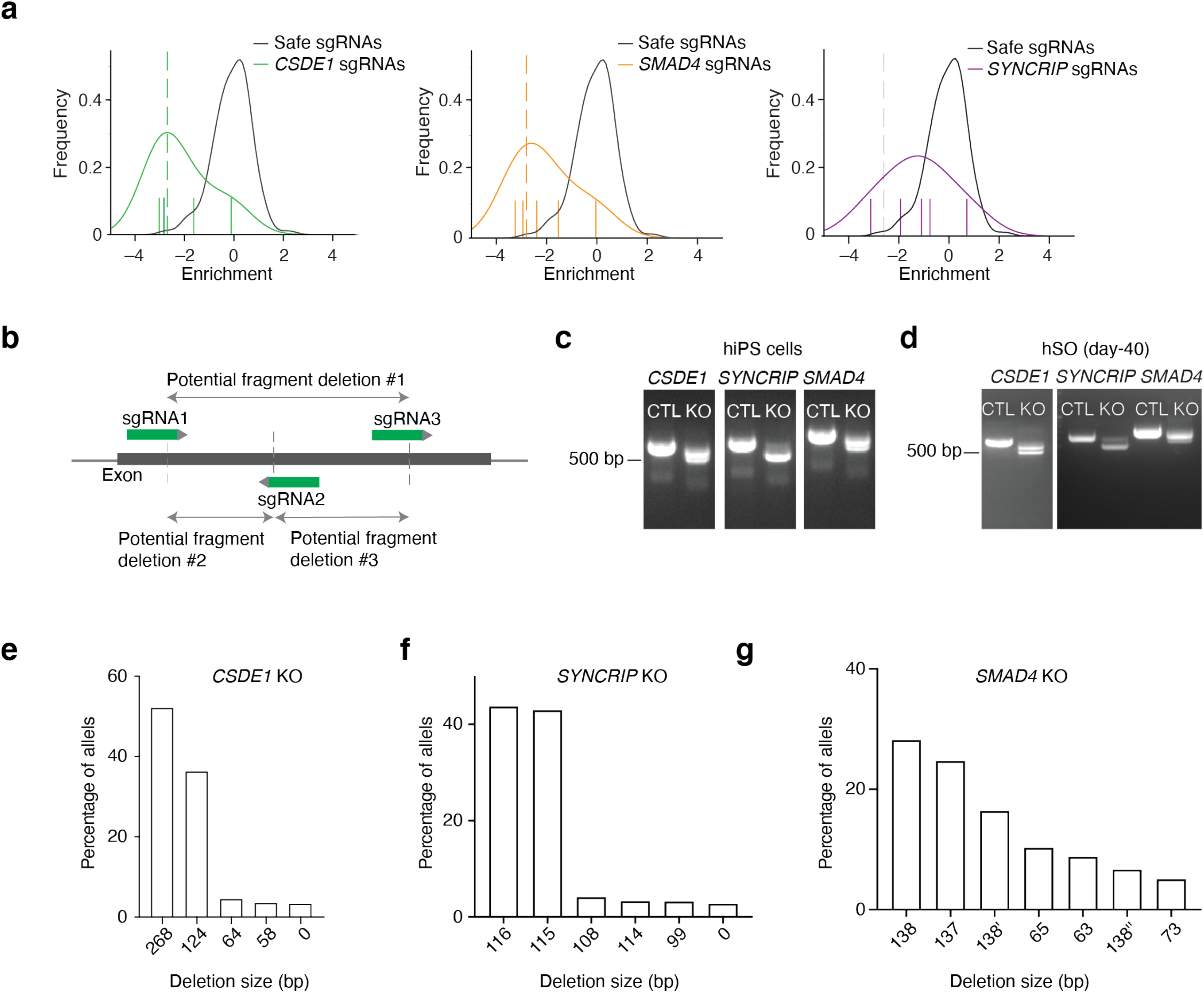
Confirmation of the KO efficiency of the selected candidate genes: *CSDE1* and *SMAD4*. **a**. Estimation of gene perturbation effect sizes. Log_2_ fold change enrichments, normalized to the median of negative ‘safe’-targeting controls, of sgRNAs between the Dlxi1/2::eGFP^+^ and Dlxi1/2::eGFP-samples are plotted in the indicated color; the distribution of safe-targeting sgRNA controls in black. The dashed line is the CasTLE-estimated maximum effect size. **b**. Schematic showing 3 sgRNAs designed to target an early exon of the gene of interest to generate multiple fragment deletions. **c–d**. Genotype analysis of genomic DNA extracted from hiPS cells (**c**) or day 40 hSO (**d**) showing the PCR amplicon of individual candidate genes from Cas9-CTL and KO cell pools. **e–g**. NGS analysis of amplicons amplified from genomic DNA extracted from day 40 hSO derived from *CSDE1* KO (**e**), *SYNCRIP* KO (**f**), and *SMAD4* KO (**g**) cells. The percentage of each allele that is more than 2% is presented. In (**g**), 138, 138’ and 138’’ indicate three different alleles that showed a 138 bp deletion.

**Supplementary Figure 5:**
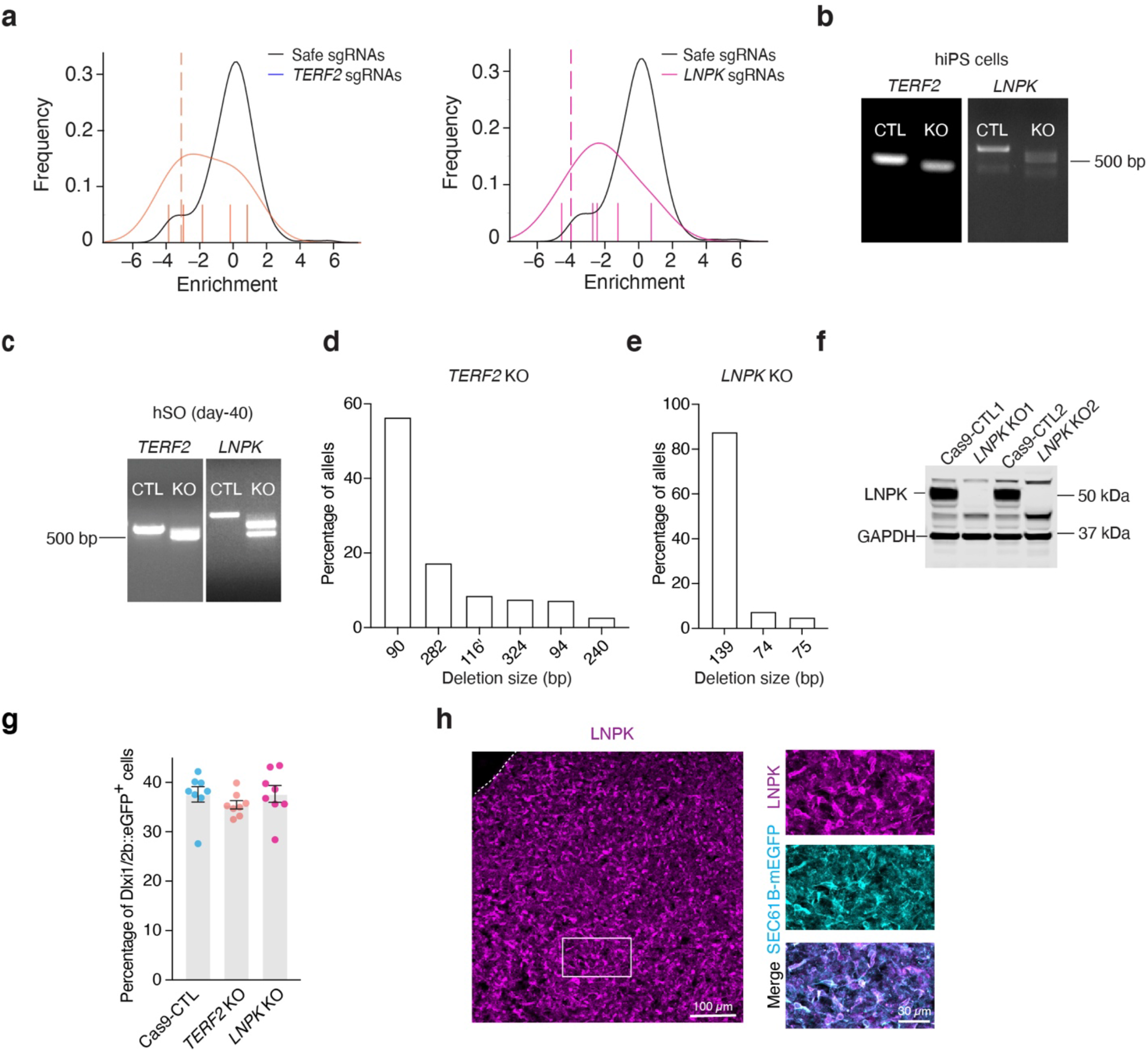
Confirmation of KO efficiency of *TERF2* and *LNPK* and immunofluorescence imaging of LNPK. **a**. Estimation of gene perturbation effect sizes. Log_2_ fold change enrichments, normalized to the median of negative ‘safe’-targeting controls, of sgRNAs between the Dlxi1/2::eGFP^+^ and Dlxi1/2::eGFP-samples are plotted in the indicated color; the distribution of safe-targeting sgRNA controls is shown in black. The dashed line is the CasTLE-estimated maximum effect size. **b–c**. Genotype analysis of genomic DNA extracted from hiPS cells (**b**) or day 40 hSO (**c**) showing the PCR amplicon of individual candidate genes from Cas9-CTL and KO cell pools. **d–e**. NGS analysis of amplicons amplified from genomic DNA extracted from day 40 hSO derived from *TERF2* KO (**d**) and *LNPK* KO (**e**) cells. The percentage of each allele that is more than 2% is shown. In (**d**), *TERF2* 116’ has an additional 21 bp insertion. **f**. Representative western blotting image showing LNPK expression. Samples are from 2 independent differentiations. GAPDH was used as a loading control. **g**, Percentage of Dlxi1/2b::GFP^+^ cells in day 57 hSO from Cas9-CTL and KO samples (n = 8 from 2 independent differentiations). Each organoid was processed as one sample. Data are presented as mean ± SEM; Kruskal-Wallis test (P = 0.1316). **h**. Left, a representative immunofluorescence image showing LNPK expression in hSO. Right, inset showing LNPK co-localizes with SEC61B-mEGFP in hSO derived from SEC61B-mEGFP hiPS cells.

**Supplementary Figure 6:**
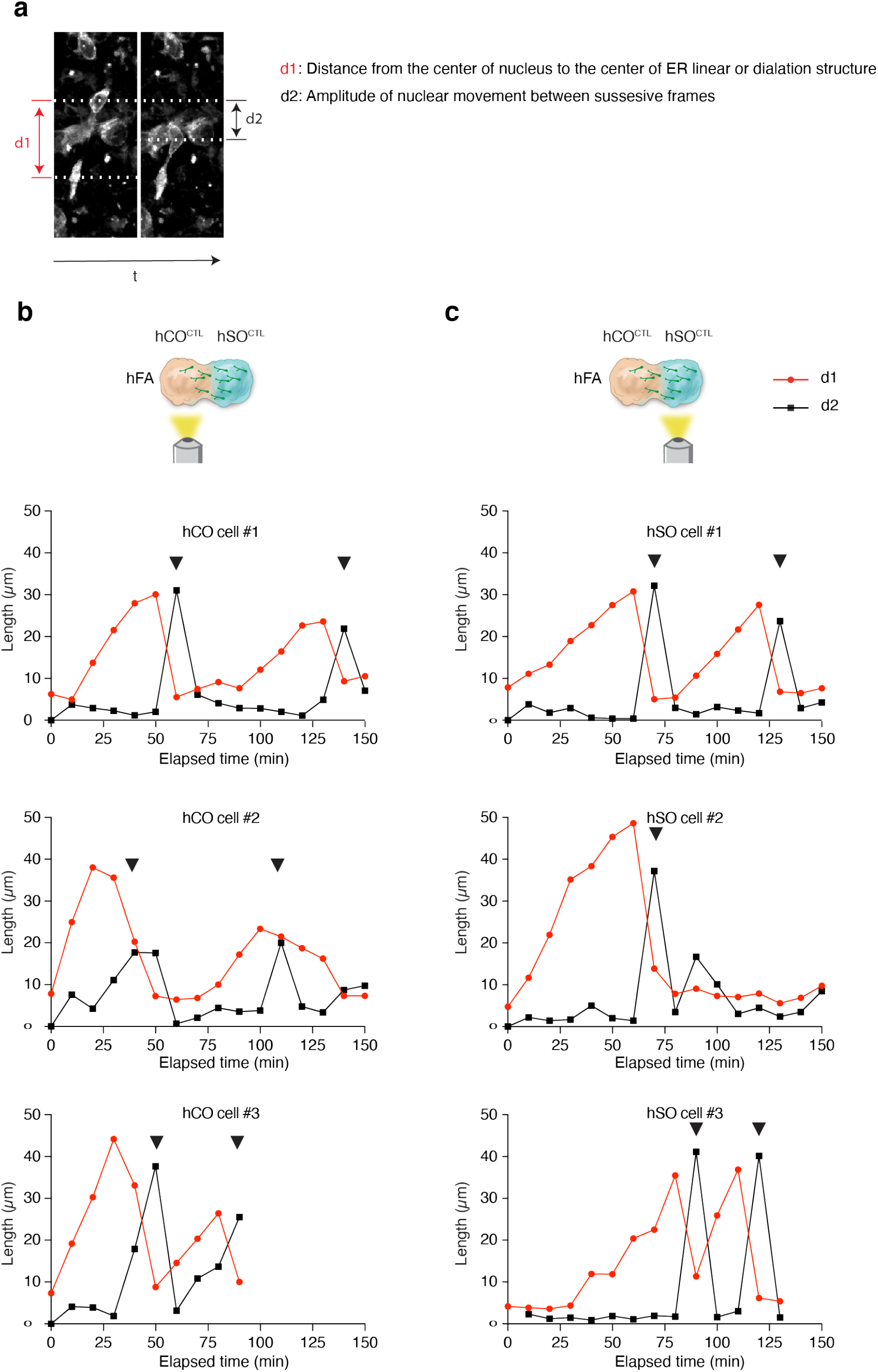
ER is displaced prior to nuclear translocation in migrating human neural cells. **a**. Example image showing measurements to study ER displacement and nuclear translocation. **b–c**. Relation of nuclear and ER movement in example migrating SEC61B-mEGFP^+^ cells in hFA (related to Figure 4c). Black triangles indicate nuclear translocations.

**Supplementary Figure 7:**
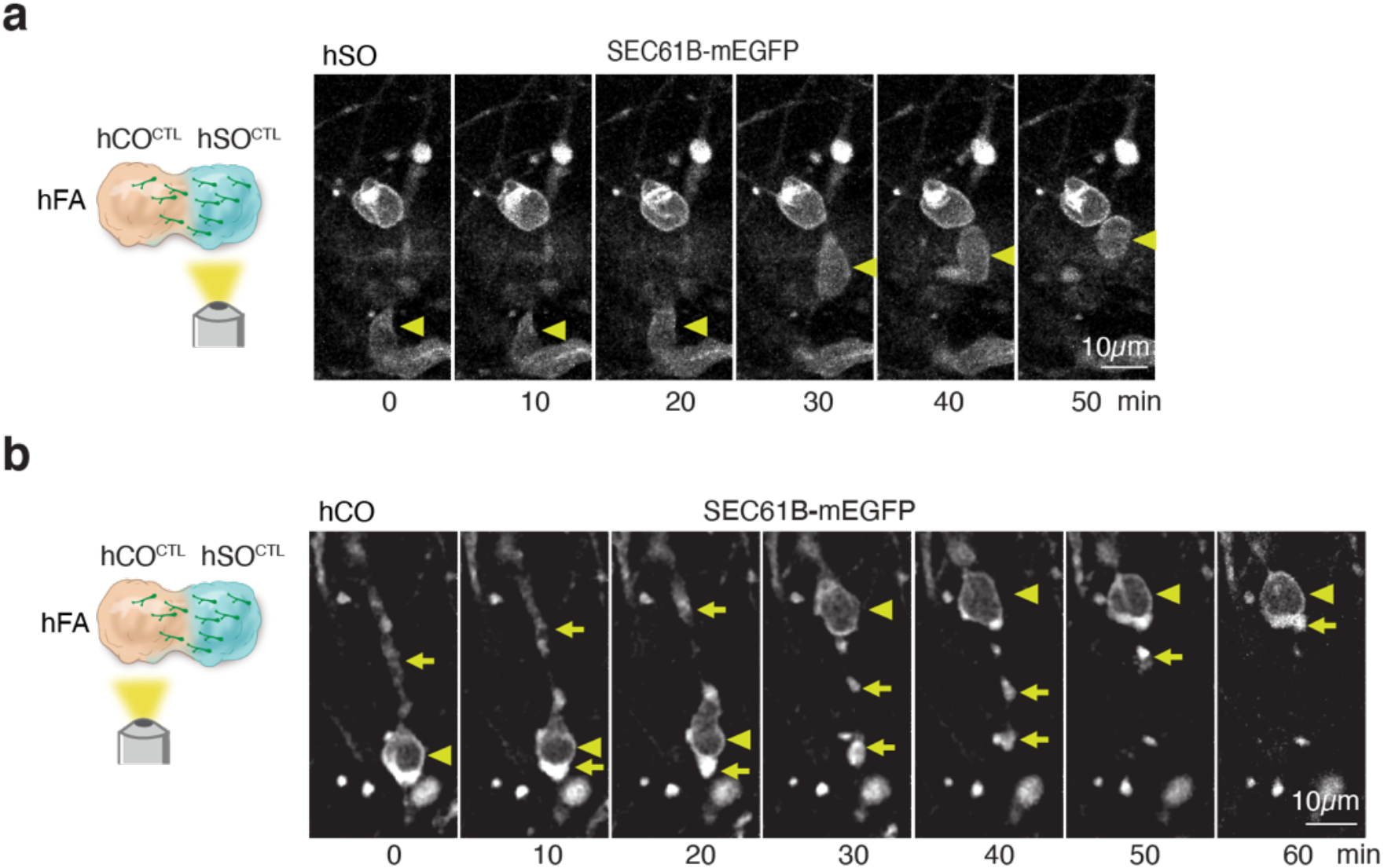
Other patterns of ER dynamic in saltatory moving cells. **a–b**. Time-lapse sequences of a SEC61B-mEGFP^+^ cell in hSO (**a**) or hCO (**b**) moving in a saltatory pattern. The hFA is generated by assembly of SEC61B-mEGFP hSO with unlabeled hCO. Triangles mark the nucleus and arrows mark ER localization at the leading branch or the trailing process during saltatory movement. The cell in (**a**) showed no detectable ER displacement. The cell in (**b**) has partial ER signal “left behind” the nucleus after nuclear translocation.

**Supplementary Figure 8:**
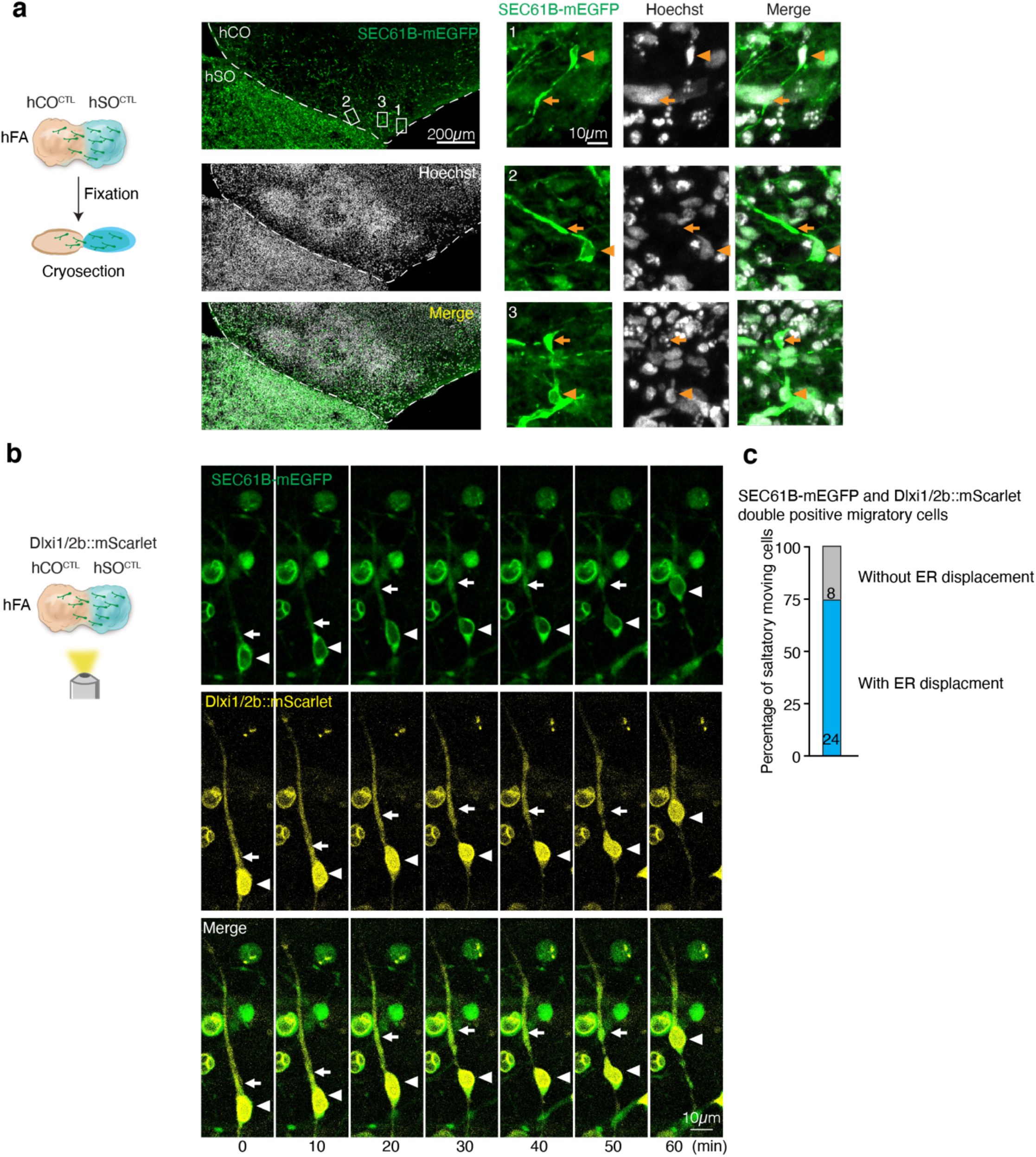
Migration-related ER displacement in fixed hFA and Dlxi1/2b::mScarlet^+^ cells. **a**. Left: Immunofluorescence cryosection images of hFA obtained by assembly of unlabeled hCO with SEC61B-mEGFP hSO. Right: Inset showing the ER distribution in SEC61B-mEGFP^+^ cells migrated into hCO. Triangles mark the nucleus and arrows mark the linear or dilated structure of the ER in the leading branch. From 2 independent experiments in n = 7 hFA. **b**. Representative time-lapse sequences showing ER displacement during the saltatory movement of a SEC61B-EGFP^+^ cell that were co-labelled with Dlxi1/2b::mScarlet (at ∼60 days post-assembly). Triangles mark the nucleus and arrows mark the linear or dilated structure of the ER during migration (n = 5 hFA). **c**. Percentage of SEC61B-mEGFP and Dlxi1/2b::mScarlet double positive migratory cells showing ER displacement. Live images were taken from both hCO and hSO sides (n = 5 hFA).

**Supplementary Figure 9:**
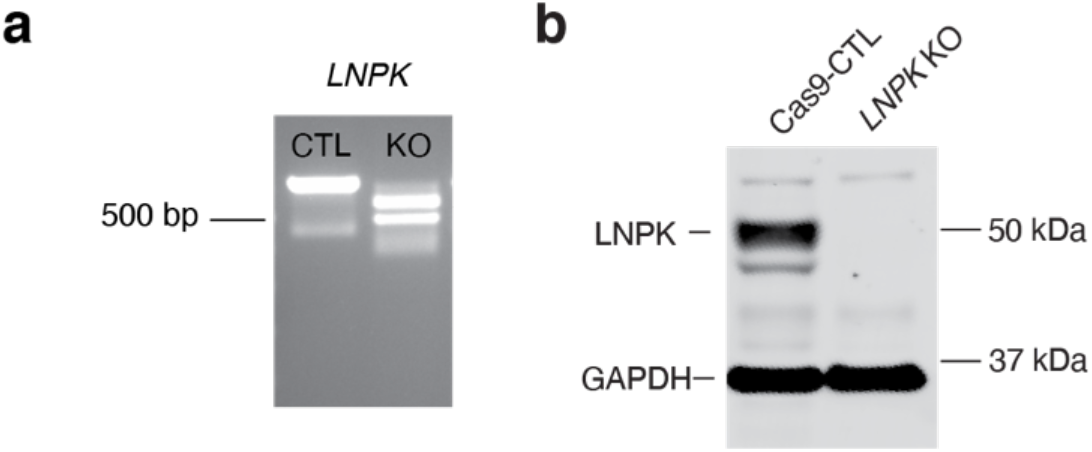
Confirmation of the deletion of LNPK in the KO cell pool engineered from the SEC61B-mEGFP hiPS cell line. **a**. Genotype analysis of genomic DNA extracted from hSO showing the PCR amplicon of *LNPK* from Cas9-CTL and *LNPK* KO cells. **b**. Representative western blotting image showing LNPK expression in *LNPK* KO and Cas9-CTL. GAPDH was used as a loading control. This experiment has been repeated 3 times.

**Supplementary Table 1: List of 611 NDDs genes**.

**Supplementary Table 2: List of genes included in the screen**.

**Supplementary Table 3: Candidate genes of the interneuron generation screen**.

**Supplementary Table 4: Candidate genes of the Interneuron migration screen**.

**Supplementary Table 5: Adjusted P values for Figure 2f**.

**Supplementary Table 6: sgRNA sequences used to generate KO cell pools**.

**Supplementary Table 7: Primers used to amplify gDNA for NGS**.

**Supplementary Table 8: Primers for RT-qPCR**.

**Supplementary Table 9: Primers for genotyping KO cells**.

**Supplementary Video 1: Z-stack images of cleared hFA and image processing to examine interneuron migration**.

**Supplementary Video 2: Live imaging showing migration of Dlxi1/2b::eGFP**^**+**^ **cells in fused hFA derived from Cas9-CTL or LNPK KO hSO fused with unlabeled hCO**.

**Supplementary Video 3: Live imaging showing a saltatory migrating SEC61B-mEGFP**^**+**^ **cell in fused hFA with ER forward migration in the leading branch prior to nuclear translocation**.

**Supplementary Video 4: Live imaging showing migration of SEC61B-mEGFP**^**+**^ **cell in fused hFA derived from Cas9-CTL or LNPK KO hSO fused with unlabeled hCO**.

